# The *Drosophila* Estrogen-Related Receptor promotes triglyceride storage within the larval fat body

**DOI:** 10.1101/2024.09.13.612925

**Authors:** Tess D. Fasteen, Melody R. Hernandez, Robert A. Policastro, Maria C. Sterrett, Gabriel E. Zenter, Jason M. Tennessen

## Abstract

The Estrogen-Related Receptor (ERR) family of nuclear receptors (NRs) serve key roles in coordinating triglyceride (TAG) accumulation with juvenile growth and development. In both insects and mammals, ERR activity promotes TAG storage during the post-embryonic growth phase, with loss-of-function mutations in mouse *Esrra* and *Drosophila melanogaster dERR* inducing a lean phenotype. However, the role of insect ERRs in controlling TAG accumulation within adipose tissue remains poorly understood, as previous transcriptomic and metabolomic studies relied on whole animal analyses. Here we address this shortcoming by using tissue-specific approaches to examine the role of dERR in regulating lipid metabolism within the *Drosophila* larval fat body. We find that dERR autonomously promotes TAG accumulation within fat body cells and regulates expression of genes involved in glycolysis, β-oxidation, and mevalonate metabolism. As an extension of these results, we not only discovered that *dERR* mutant fat bodies exhibit decreased expression of known dHNF4 target genes but also found that dHNF4 activity is decreased in *dERR* mutants. Overall, our findings indicate that dERR plays a multifaceted role in the larval fat body to coordinate lipid storage with developmental growth and hint at a conserved mechanism by which ERR and HNF4 homologs coordinately regulate metabolic gene expression.

## INTRODUCTION

Growth and development must be closely coordinated with metabolism to fulfill the differential energetic requirements of each life stage (Garfinkel et al., 2024, Miyazawa and Aulehla, 2018). Among the key metabolites that dictate both growth rate and developmental progression are lipids, which serve in developmental signaling pathways, act as building blocks for membranes and other cellular structures, and function as energetic reservoirs that support major developmental events and ensure survival during bouts of stress and starvation (Palm and Rodenfels, 2020, Zhu and Han, 2014, Watts and Ristow, 2017, Ho et al., 2004, Srinivasan, 2015). As a result, disruptions in lipid metabolism can significantly alter growth trajectories, adult lifespan, and reproductive capacity. For example, in humans, body fat percentage is more closely linked with the onset of mammalian puberty than chronological age (Baker, 1985, Huang et al., 2023, Li et al., 2022). Similarly, maternal obesity disrupts metabolic cues during fetal development, resulting in altered hormone levels, decreased adipogenesis, induction of lipolysis, and precocious onset of puberty in the offspring (Scheidl et al., 2023, Barrea et al., 2022). Abnormal lipid metabolism, however, not only influences developmental progression, but also results in increased risk of developing metabolic disorders later in life, including type 2 diabetes, heart disease, strokes, osteoarthritis, and other comorbidities (Marcus et al., 2022, Menendez et al., 2022, Palacios-Marin et al., 2023). Thus, the molecular mechanisms that coordinate lipid metabolism with growth and development broadly influence animal health and disease progression.

The fruit fly *Drosophila melanogaster* has emerged as a powerful genetic system for studying the coordinate regulation of lipid metabolism and juvenile growth (Heier et al., 2021, Lehmann, 2018, Gáliková and Klepsatel, 2023, Musselman and Kühnlein, 2018). During the course of larval (juvenile) development, the fly accumulates large TAG reserves that serve diverse and essential biosynthetic and energetic functions (Carvalho et al., 2012, Heier et al., 2021, Church and Robertson, 1966). Not only do these TAG stores buffer larval development against bouts of nutrient deprivation (Rodenfels et al., 2014, Bi et al., 2012), but larval fat tissue also persists throughout metamorphosis into the early adult stages, where TAG supports the energetic and biosynthetic requirements of newly eclosed adults (Carvalho et al., 2012, Aguila et al., 2007, Hofbauer et al., 2021). Moreover, TAG serves as a carbon sink during periods of nutrient excess, preventing sugar and lipid intermediates from accumulating to levels that induce metabolic disease phenotypes (Musselman et al., 2013, Garrido et al., 2015).

The insect fat body is one of the key tissues responsible for coordinating insect TAG metabolism with larval growth and development. In its role as the functional equivalent of both mammalian liver and adipose tissue, the fat body monitors the systemic levels of lipids and other metabolites, controls the balance between macromolecular storage and interorgan metabolite transport, and produces secreted factors that regulate systemic growth (Li et al., 2019, Skowronek et al., 2021, Texada et al., 2020, Danielsen et al., 2013, Tennessen and Thummel, 2011). Considering that many of the nutrient sensors and signal transduction cascades that regulate metabolism within the larval fat body are highly conserved, this organ represents a powerful model for studying how metabolism, and lipid metabolism in particular, is regulated in the context of animal development.

Among the conserved factors that function within the fat body, dERR, the sole *Drosophila* ortholog of the ERR family of nuclear receptors, stands out as a promising candidate for examining how lipid metabolism and developmental progression are coordinately regulated. Although insect ERRs primarily regulate genes involved in carbohydrate metabolism (Long et al., 2020, Geng et al., 2024, Tennessen et al., 2011, Beebe et al., 2020), studies in both *Drosophila melanogaster* and the mosquito *Aedes aegypti* demonstrate that loss of ERR activity results in a lean phenotype (Geng et al., 2024, Tennessen et al., 2011, Beebe et al., 2020). Similarly, while *Esrra* mutant mice exhibit relatively normal development, these animals are lean, exhibit defects in lipid uptake, and are resistant to diet-induced obesity (Luo et al., 2003, Carrier et al., 2004). Such findings not only indicate that the ERR family members play an ancient and conserved role in coordinating lipid metabolism with juvenile development, but also position *Drosophila melanogaster* as an ideal model for understanding how this NR family member functions within adipose tissue. However, previous studies that examined the role of dERR in regulating lipid metabolism largely relied on whole animal multi-omic assays and thus overlooked fat body-specific dERR functions (Geng et al., 2024, Tennessen et al., 2011, Beebe et al., 2020). Here, we address this issue by examining how dERR influences lipid metabolism specifically within the larval fat body.

Using a combination of quantitative lipidomics and tissue-specific genetic manipulations, we demonstrate that dERR activity within the larval fat body is sufficient to promote TAG storage. When dERR is absent from this tissue, larvae exhibit decreased levels of both TAGs and acylcarnitines (ACARs) and display a lean phenotype. Moreover, RNA-seq analysis of fat bodies isolated from *dERR* mutant larvae reveals a significant decrease in not only glycolytic gene expression, but also the misexpression of an array of genes involved in both lipid synthesis and lipid catabolism. Intriguingly, this RNA-seq dataset demonstrated that loss of dERR activity results in a significant down-regulation of genes associated with fatty acid β-oxidation – a result that has not been previously observed in whole animal RNA-seq studies. Finally, we demonstrate that the nuclear receptor dHNF4 displays decreased activity in *dERR* mutants. Considering that dHNF4 is a key regulator of fatty acid β-oxidation in fly larvae (Palanker et al., 2009), our findings inform a model in which these highly conserved nuclear receptors function in the fat body to coordinately regulate aspects of lipid metabolism within the context of larval growth.

## RESULTS

### Lipidomic analysis of *dERR* mutants reveals significant reduction of TAG levels

To determine how dERR promotes larval lipid accumulation, we used a quantitative lipidomics approach to measure the concentration of 537 lipid species in both *dERR^1/2^* mutants and *dERR^1/+^* heterozygous controls (Table S1). Principal component and differential expression analyses of the resulting datasets revealed significant differences between control and mutant larvae (Figure S1), with a total of 17 lipid species significantly changing in abundance (Log2 fold change ≥ 1; adjusted *p* < 0.05; Figure 1A, Table S2). Among these lipids, the most significantly altered species are triglycerides (TAGs; Figure 1A,B), with six individual TAG molecules meeting or exceeding the cutoff threshold. Five of these six lipids rank among the most abundant TAG species present within larvae (Table 1), and subsequent analyses revealed that nine of the 25 most abundant TAGs are significantly decreased in *dERR* mutants relative to heterozygous controls. Notably, the two TAGs with the highest concentration in control larvae, TG_16:0_16:1_18:1 and TG_14:0_16:1_18:1, are decreased ∼40% in mutant larvae relative to the controls (Table 1, Figure 1B,C; q>0.01). Together, these observations demonstrate that loss of dERR activity results in a significant decrease of TAG stores.

**Figure 1.**
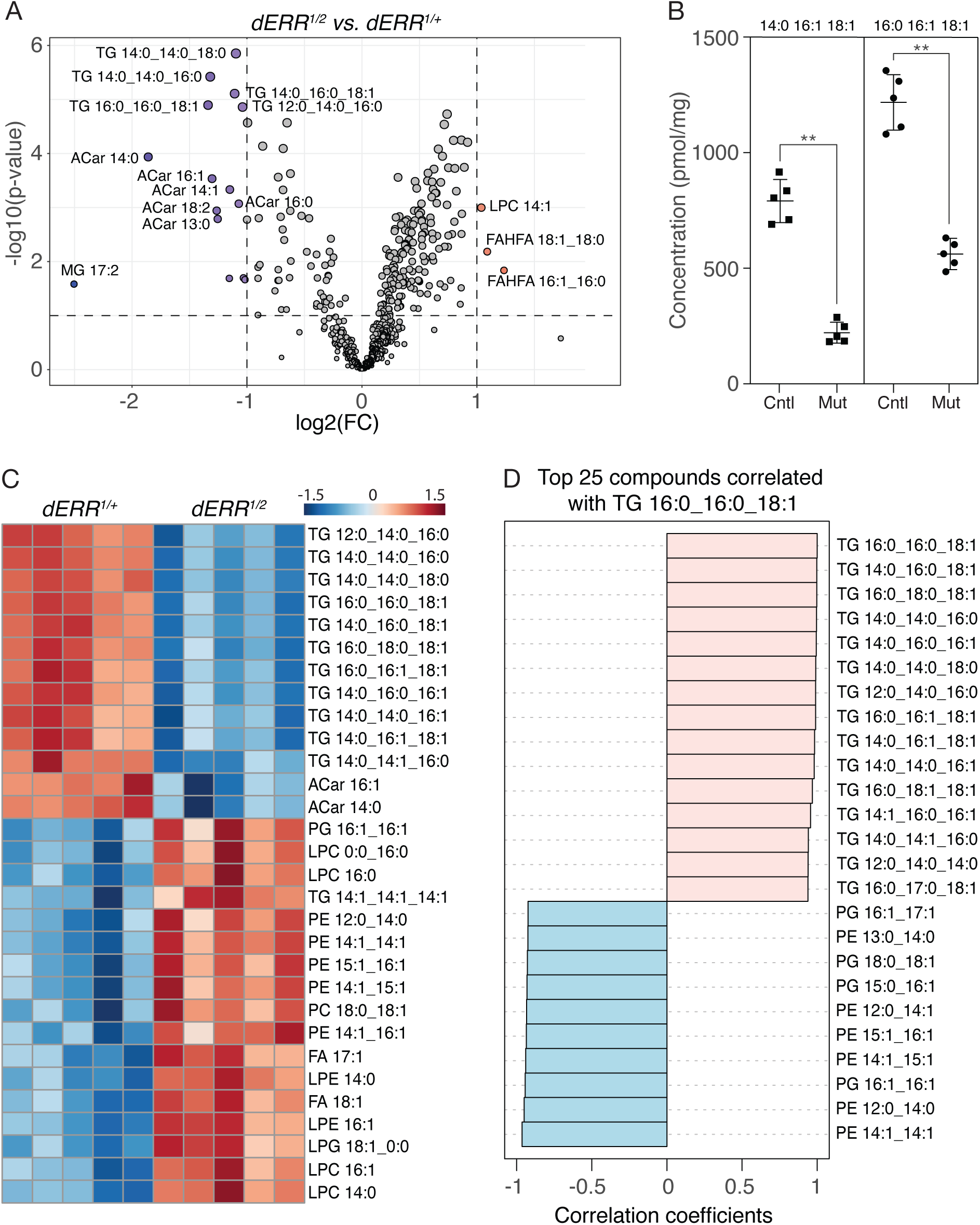
Quantitative lipidomic analysis of *dERR* mutant larvae. *dERR^1/2^*mutant larvae and *dERR^1/+^* controls were analyzed using a quantitative lipidomic approach that measured the concentration of 537 lipid species. (A) Differences in lipid abundance between the mutant and control groups are represented with a volcano plot. Dashed vertical line represents a log_2_ fold change (FC) of 2, and dashed horizontal line represents *P*=0.1. (B) Abundance of the triglycerides 16:0_16:1_18:1 and 14:0_16:1_18:1 in *dERR* mutant larvae (Mut) as compared to controls (Cntl). (C) The top 30 significantly altered lipid species between mutant and control larvae are represented in a heat map and ordered based on *P* value calculated using a two sample t-test. (D) The top 25 compounds correlated with TAG 16:0_16:0_18:1. n=5 biological replicates for each genotype and each replicate consists of 20 mid-L2 larvae. \*\**P* <0.01. *P-*value calculated using a Kolmogorov-Smirnov test. All panels were generated using Metaboanalyst 6.0.

**Table 1.**
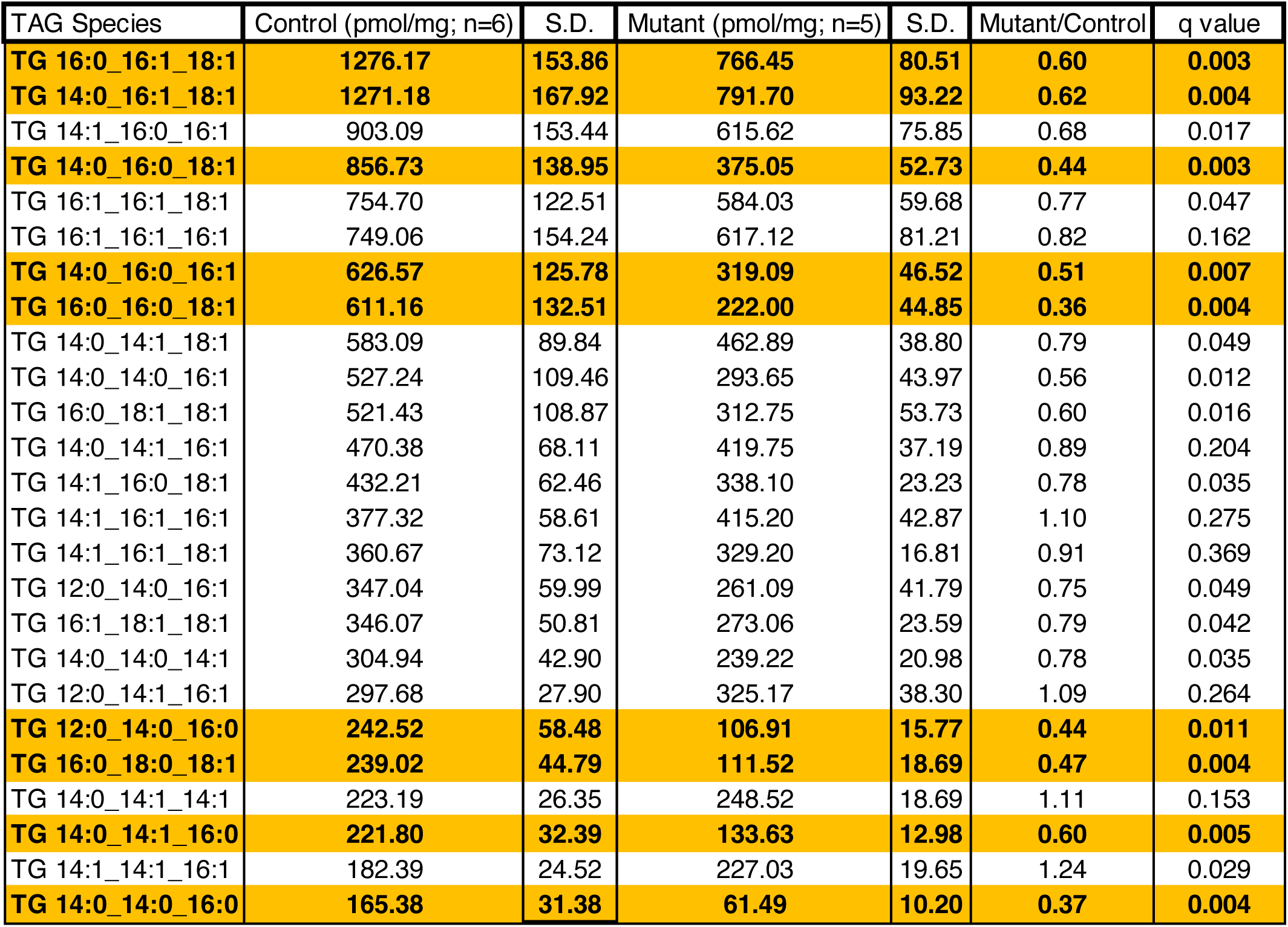
Quantification of the 25 most abundant TAG species in ERR^1/2^ mutants and ERR^1/+^ controls. q<0.01 for highlighted rows. Standard deviation is abbreviated S.D.

Beyond the observed changes in TAG abundance, the levels of several other lipids are significantly altered in *dERR* mutants when compared with control. Notably, ACAR species were down-regulated greater than 2-fold in mutants when compared with the heterozygous control (Table S2, Figure 1A), raising the possibility that fatty acid β-oxidation is decreased in *dERR* mutants. Moreover, when we reanalyzed the lipidomic data using a less stringent cutoff (absolute FC>1.5; adjusted *P*<0.05; Table S3), several phospholipids (phosphatidylcholine [PCs]; phosphatidylethanolamine [PEs]; phosphatidylglycerols [PGs]) and lysophospholipids ([LPC]; [LPG]; [LPE]) were found to be elevated in *dERR* mutants (Figure 1C). In fact, a correlation analysis of TG_16:0_16:1_18:1 revealed a significant inverse correlation between PE levels and TAG levels in *dERR* mutant larvae (Figure 1D). Overall, our analysis reveals that of those lipids significantly altered in *dERR* mutants, there are significant decreases in stored lipids and acylcarnitines as well as a subtle increase in phospholipids.

### dERR expression within the fat body promotes TAG accumulation

Previous studies that examined the influence of dERR on larval TAG metabolism relied on measurements from whole animal homogenates. To better understand the tissue-specific mechanisms by which ERR regulates TAG accumulation, we focused on the fat body, which is the primary TAG storage site. As an initial approach to determine whether the global decrease in *dERR* mutant TAG levels was reflected in the fat body, we stained mutant and control larvae using the lipophilic dyes Solvent Black 3 (SB3) and Nile Red. When compared with controls, *dERR* mutant fat bodies exhibited significantly lower levels of both SB3 and Nile Red staining (Figure 2A-D), indicating that loss of dERR activity results in decreased fat body TAG accumulation.

**Figure 2:**
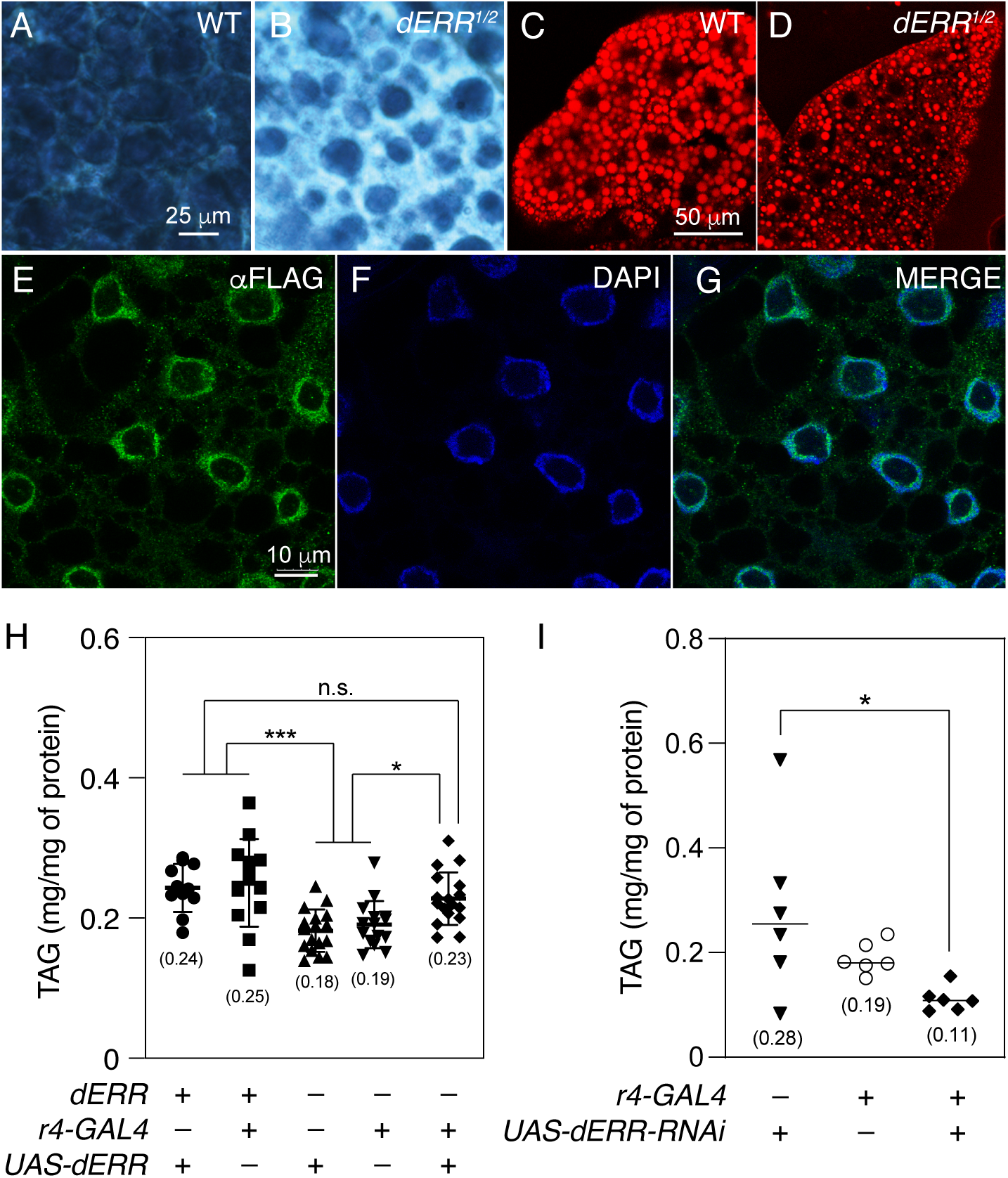
dERR promotes TAG accumulation in the larval fat body. (A-D) Fat bodies from *w^1118^* controls and *dERR^1/2^* mutants were dissected from mid-L2 larvae (∼60 hrs after egg-laying), fixed, and stained with either (A,B) Solvent Black 3 or (C,D) Nile Red. Scale bar in (A) also applies to (B). Scale bar in (C) also applies to (D). (E-F) Fat bodies from mid-L2 larvae expressing a previously dERR-GFP-StrepII-Flag transgene were stained with αFlag antibody and DAPI. Scale bar in panel (E) applies to (F) and (G). (H) TAG levels were quantified relative to soluble protein in whole body extracts from heterozygous control strains *r4-Gal4/+* and *+/UAS-ERR*, the *dERR* mutant strains *r4-Gal4/+*; *dERR^1/2^* and *+/UAS-ERR*; *dERR^1/2^*, and the rescue strain *r4-Gal4 +/+ UAS-dERR*; *dERR^1/2^*. (I) TAG levels were quantified relative to soluble protein in whole body extracts from heterozygous control strains *r4-Gal4/+* and *+/UAS-dERR-RNAi*, as well as larvae expressing the *UAS-dERR-RNAi* transgene under the control of *r4-Gal4*. Data analyzed using an ordinary ANOVA test followed by a Holm-Sidak test for multiple comparisons. * *P*<0.05. \*\*\**P*<0.001.

The SB3 and Nile Red staining results raised the question as to whether dERR functions autonomously within fat body cells to promote TAG accumulation. To test this possibility, we first examined if dERR is expressed L2 larval fat bodies using a previously described transgene that expresses a *dERR-FLAG-GFP* fusion protein from the endogenous *dERR* promoter (Venken et al., 2009). Our analysis revealed that dERR is highly expressed within larval fat bodies and primarily localized to the nucleus, although we also noted low levels of dERR present within the cytosol (Figure 2E-H). This result, together with previous observations demonstrating that a dERR ligand-binding domain reporter is active within larval fat body cells (Palanker et al., 2006), suggested that *dERR* mutants exhibit decreased TAG levels due to loss of dERR expression within this tissue. We tested this possibility by using the fat body-specific driver *r4-Gal4* to (i) express a *UAS-dERR* transgene in a *dERR* mutant background, and (ii) express an *UAS-ERR-RNAi* transgene in the wild-type fat body. Consistent with our hypothesis, expression of dERR only in fat body cells is sufficient to restore normal TAG levels in *dERR* mutants (Figure 2H). Similarly, we observed a decrease in the TAG levels of *r4-ERR-RNAi* larvae when compared with the control strains (Figure 2I). In this regard, we would note that although *P>0.05* when comparing TAG levels between the *UAS-ERR-RNAi* control strain and *r4-ERR-RNAi* larvae, TAG concentrations in *r4-ERR-RNAi* samples were 50% lower that *UAS-ERR-RNAi* control samples (Figure 2I). Together, these observations indicate that dERR acts within larval fat body cells to promote normal TAG homeostasis and align with similar observations in adult fat body cells.

### Genes involved in carbohydrate metabolism and β-oxidation exhibit decreased expression in *dERR* mutant fat bodies

To better understand the underlying metabolic mechanisms leading to decreased TAG levels within *dERR* mutant fat bodies, we used RNA-seq to analyze gene expression in (i) the fat bodies dissected from *dERR^1/2^* mutants as well as *dERR^1/+^* heterozygous controls, and (ii) whole larvae of the same genotypes (Tables S4, S5). These analyses revealed gene expression changes within the fat body that were unique compared to those observed in the whole animal analysis; only 40 genes exhibited significantly altered expression levels in both datasets (Figure 3A, Table S6 and S7), with most of these genes encoding enzymes involved in glycolysis and carbohydrate metabolism (Table S6). In addition, principal component analysis highlighted that the tissue type segregated along the PC1 axis, which accounts for over 84% of the variance (Figure 3B), while genotype separated along PC2 and accounted for less that 5% of the variance (Figure 3B). This supports the conclusion that the *dERR* mutant gene expression profile within the fat body significantly differs from that of the whole animal.

**Figure 3.**
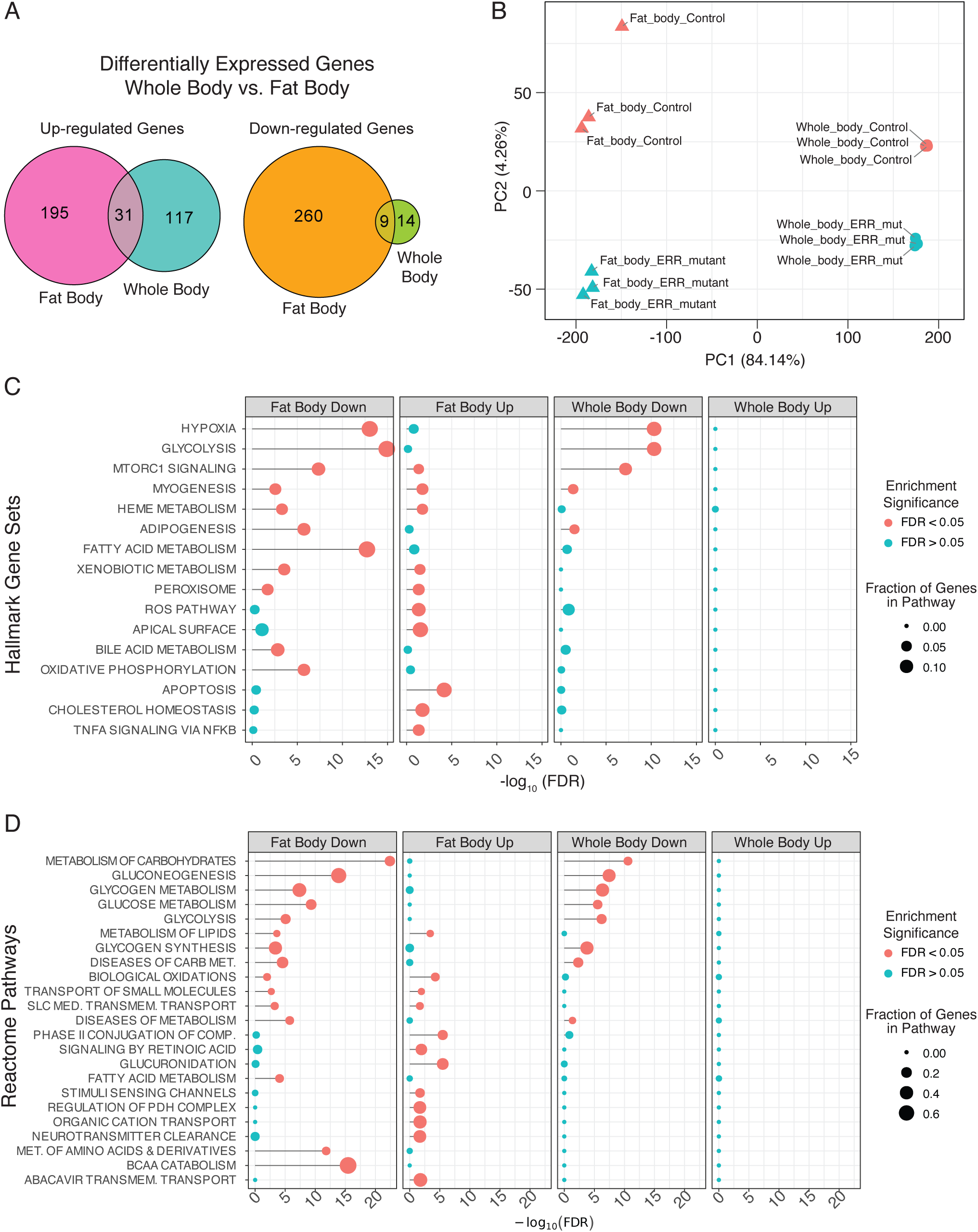
Tissue-specific RNA-seq analysis of *dERR* mutants compared with heterozygous controls. RNA isolated from whole mid-L2 *dERR^1/2^* mutant larvae and *dERR^1/+^* controls, as well as from dissected fat bodies from those two genotypes, were compared using RNA-seq. (A) Euler plot indicating the number of genes that were either down-regulated or up-regulated in both *dERR* mutant fat bodies and whole animals. (B) PCA plot of the RNA-seq samples. Samples separate along the PC1 axis based on tissue and the PC2 axis according to genotype. (C,D) Term enrichment analysis of the RNAseq data was conducted using the (C) “Hallmark Gene Sets” and (D) “Reactome Pathways” collections in the Molecular Signatures Database (MSigDB). Data from the whole body and fat body were analyzed separately as labeled.

To better understand how dERR influences gene expression, we conducted a series of Gene Ontology (GO) analyses of the whole body and fat body RNA-seq datasets. Using the Hallmark Gene Sets and Reactome Pathway databases, GO analysis revealed whole body enrichment for genes associated with metabolic processes, with the most significantly overrepresented GO categories associated with carbohydrate metabolism (*e.g.,* Hypoxia, Glycolysis, Metabolism of Carbohydrates) among the downregulated genes (Figure 3CD, third panel). In the fat body dataset, however, there is additionally significant enrichment for Fatty Acid Metabolism, Adipogenesis (Figure 3C) and Metabolism of Lipids (Figure 3D). PANGEA analysis revealed a similar enrichment for GO categories associated with metabolism, with the RNAseq data generated from the *dERR* mutant fat body exhibiting significant enrichments for genes involved in both carbohydrate and lipid metabolic processes (Table S8,S9).

A closer examination of the GO categories that are significantly enriched in *dERR* mutant fat bodies revealed disruption of three metabolic pathways. As expected, genes encoding every enzymatic step in glycolysis were significantly down-regulated (Figure 4A; Table S4, S5). However, we also unexpectedly observed decreased expression of genes associated with fatty acid β-oxidation (Figure 4B; Table S4, S5), including the rate-limiting transporter *CPT1* (*whd*, FBgn0261862) and five genes that encode enzymes involved in the four downstream catabolic reactions (Figure 4B; Table S4, S5). None of these genes, however, exhibited significant expression changes in the *dERR* mutant whole animal RNA-seq dataset. We also noted that three genes associated with isoprenoid synthesis (*Hmgs*, *Hmgcr*, and *Fpps*) exhibited increased expression levels in *dERR* mutant fat bodies but were unchanged when measured in whole animal extracts (Figure 4C; Table S4, S5). Moreover, expression of a fourth gene involved in isoprenoid metabolism, *qless*, displayed opposite changes in the two datasets, with *qless* expression being increased *dERR* mutant fat bodies and decreased in the whole-body data (Figure 4C; Table S4, S5). Overall, these results suggest that dERR serves previously undescribed functions within the larval fat body to dictate fatty acid catabolism and isoprenoid synthesis.

**Figure 4.**
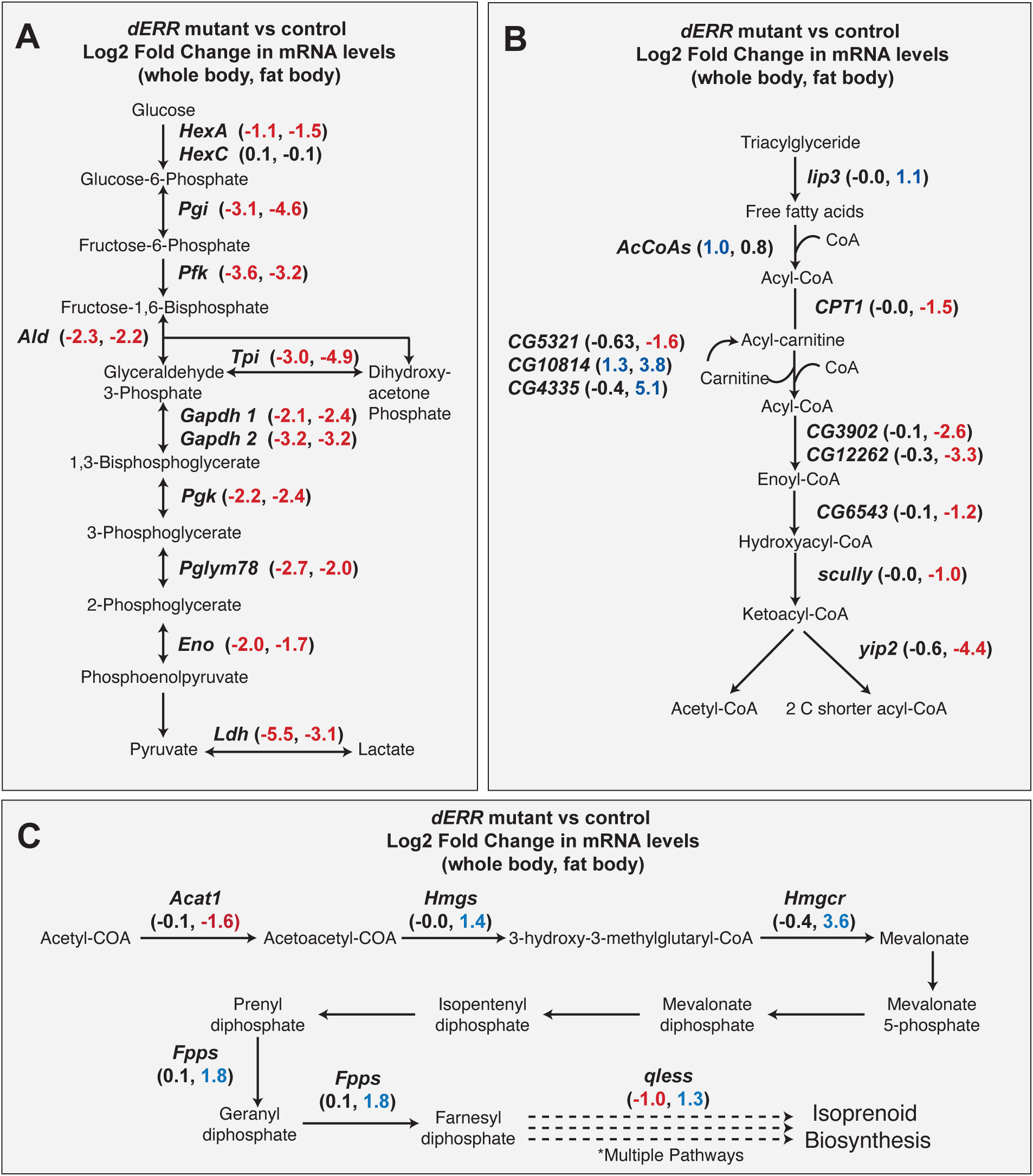
Expression of enzymes in glycolysis and β-oxidation are reduced in *dERR* mutants. Schematic diagrams illustrating how loss of dERR activity alters the expression of genes involved in (A) glycolysis, (B) β-oxidation, and (C) isoprenoid metabolism. Changes in gene expression are listed at each enzymatic step, with the first number in parenthesis indicating the Log2-fold change in whole animal extract and the second number indicating the Log2-fold change in only larval fat body. Log2 ≥ 1 represented in blue and Log2 ≤ 1 represented in red for both panels.

### HNF4 activity is decreased in *dERR* mutant fat bodies

Our RNA-seq analysis indicates that dERR can influence the expression of genes associated with fatty acid β-oxidation. Considering that all of the β-oxidation-related genes down-regulated in *dERR* mutants are known targets of HNF4 (Palanker et al., 2009), and a recent study in mouse liver revealed an interaction between ERRα and HNF4α (Scholtes et al., 2024), our findings raise the possibility that dHNF4 activity is altered in *dERR* mutant fat bodies. We tested this hypothesis by placing a dHNF4 ligand binding domain (LBD) reporter (*hs-Gal4-HNF4-LBD, UAS-nLacZ*) into a *dERR* mutant background and stained for β-galactosidase activity. This HNF4-LBD reporter has previously been shown to be active at low levels in the larval fat body under fed conditions. Consistent with these prior studies, the HNF4-LBD reporter was active at detectable but low levels in the fat body of the *dERR^1/+^*heterozygous controls (Figure 5B), however, reporter activity was undetectable in the *dERR^1/2^* mutant background (Figure 5C). In fact, the amount of lacZ staining in *hs-Gal4-HNF4-LBD UAS-nLacZ; dERR^1/2^* larvae (Figure 5C) was indistinguishable from the *UAS-nLacZ* control strain (Figure 5A), thus indicating that HNF4 activity is reduced in *dERR* mutants.

**Figure 5.**
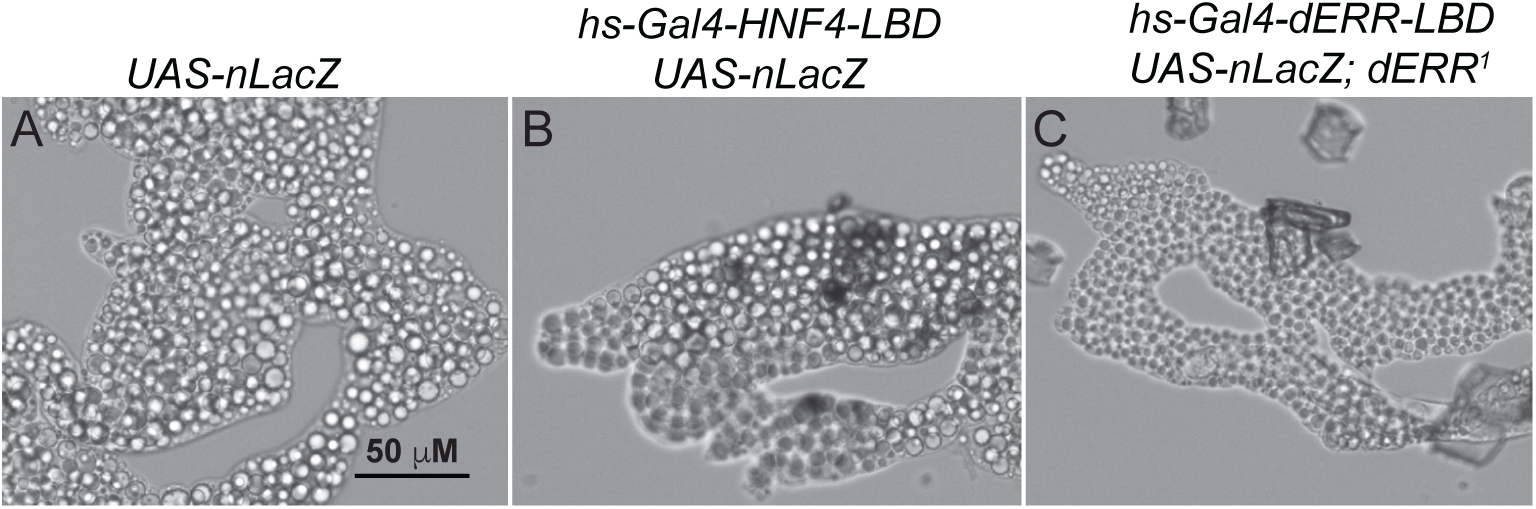
dHNF4 activity reporter in a *dERR* mutant background. HNF4 activity was visualized via X-Gal staining using *hs-Gal4-dHNF4-LBD* and *UAS-nLacZ* transgenes that were placed in a *dERR* mutant genetic background. Figure shows staining for (A) *UAS-nLacZ* control strain, (B) *hs-Gal4-dHNF4-LBD, UAS-nLacZ; dERR^1/+^* heterozygous control, and (C) *hs-Gal4-dHNF4-LBD, UAS-nLacZ; dERR^1^* mutants.

## DISCUSSION

While loss-of-function mutations in both mouse *Esrra* and insect *ERR* genes are well-documented to induce a lean phenotype (Luo et al., 2003, Beebe et al., 2020, Geng et al., 2024), the mechanisms by which these ERR family members function in adipose tissue to regulate TAG storage remain poorly understood. Here, we addressed this knowledge gap by conducting RNA-seq analysis of fat bodies isolated from *Drosophila melanogaster ERR* mutants. Our studies revealed that fat bodies from *dERR* mutants exhibit significant decreases in the expression of genes involved in both carbohydrate and lipid catabolism. This finding was unexpected, as previous studies of dERR in both larvae and adults failed to identify a link between dERR activity and the expression of enzymes involved in lipid catabolism (Beebe et al., 2020, Tennessen et al., 2011). Considering that our lipidomic studies revealed significant decreases in both TAG and ACAR concentrations, our findings suggest that dERR can influence the expression of enzymes involved in lipid metabolism.

These results raise the question as to how dERR regulates lipid metabolism in the fat body. While previous ChIP-seq analysis of adult flies failed to identify direct binding of dERR to any genes involved in lipid synthesis or β-oxidation (Beebe et al., 2020), this analysis relied on chromatin isolated from whole animals and would likely overlook fat body-specific dERR target genes. Consistent with this possibility, a recent study in *A. aegypti* demonstrated that mosquito ERR can bind to the promoter region of *fatty acid synthase* (Geng et al., 2024), thus suggesting that insect ERRs can directly regulate some genes involved in lipid metabolism. Future studies in *Drosophila melanogaster* should focus on identifying the dERR target genes specifically within the fat body to determine what aspects of lipid metabolism are directly regulated by dERR activity.

Our RNA-seq analysis also unexpectedly revealed increased expression of enzymes involved in isoprenoid metabolism within *dERR* mutant fat bodies. Previous studies in *Drosophila* have implicated these enzymes in germ cell migration, juvenile hormone production, heart development, and signal transduction (Yi et al., 2006, Deshpande et al., 2013, Santos and Lehmann, 2004, Barton et al., 2024, Bellés et al., 2005, Deshpande and Schedl, 2005, Deshpande et al., 2009). While we unsure as to why isoprenoid metabolism appears elevated in *dERR* mutants, we would note that *Drosophila* hedgehog signaling is dependent on this pathway (Danielsen et al., 2013, Deshpande et al., 2009, Deshpande and Schedl, 2005). Since elevated hedgehog signaling is associated with decreased TAG accumulation (Suh et al., 2006, Zhang et al., 2020), perhaps increased expression of these enzymes is related to a metabolic feedback mechanism involving hedgehog signaling. Alternatively, isoprenoid metabolism also produces coenzyme Q_10_ (also known as ubiquinone) synthesis and the *Drosophila* gene *qless* is essential for Q_10_ synthesis (Grant et al., 2010). The observed increase of *qless* expression in *dERR* mutant fat bodies could therefore be related to production of a molecule in mitochondrial metabolism.

Beyond the potential that dERR acts within the fat body to directly regulate genes involved in fatty acid and isoprenoid metabolism, our analysis also suggests that changes in dHNF4 activity contribute to *dERR* mutant phenotypes. Not only does dHNF4 regulate expression of many of the β-oxidation genes that are down-regulated in *dERR* mutant fat bodies (Palanker et al., 2009), but activity of the HNF4-LBD reporter is also decreased in the fat body of *dERR* mutants. Moreover, mouse HNF4a is required for hepatic ACAR synthesis (Simcox et al., 2017) – a finding consistent with our lipidomic results which reveal decreased abundance of ACARs in *dERR* mutants. This putative interaction between dERR and dHNF4 could indicate two distinct mechanisms by which these factors coordinate lipid metabolism with larval growth and development. First, ERRa and HNF4a were recently found to bind a common set of genes in mouse liver (Scholtes et al., 2024), suggesting that dERR and dHNF4 might interact in a similar manner within the larval fat body. A second potential mechanism stems from the inability of dERR mutants to properly metabolize carbohydrates. The dHNF4 LBD is activated by free fatty acids (Palanker et al., 2009), and since carbon flux from sugars to fatty acid synthesis is likely disrupted in *dERR* mutants, decreased free fatty acid abundance in *dERR* mutants could result in decreased dHNF4 activity. We would note, however, that free fatty acids are not decreased in the *dERR* mutants, thus if this mechanism is responsible for the change in dHNF4 activity, it would require fat body-specific depletion of fatty acids. Future studies will need to evaluate both possibilities to better understand how these NRs interact within the larval fat body. Regardless of the mechanism, our findings hint at a novel tissue-specific role for dERR in the modulation of insect lipid metabolism.

## METHODS

### Drosophila Husbandry and Genetics

*Drosophila* stocks were maintained on Bloomington Stock Center medium at 25°C. For all experiments, *w^1118^; ERR^1^/TM3, P{w[+mC]=GAL4-twi.G}2.3, P{UAS-2xEGFP}AH2.3, Sb^1^ Ser^1^* (RRID:BDSC_83688) virgin females were crossed with either *w^1118^* males or w^1118^; ERR^2^/TM3, P{Dfd-GMR-nvYFP}3, Sb^1^ males (RRID:BDSC_83689) and the offspring were collected on molasses plates covered with yeast paste as previously described (Li and Tennessen, 2017). The resulting larvae were raised on the collection plates for 60 hrs at 25°C. L2 larvae of the genotypes *w^1118^; ERR^1/2^* and *w^1118^; ERR^1/+^* were collected based on the absence of GFP and YFP expression. For all genetic experiments, *UAS-transgene* expression was driven using *y*^1^ *w*; P{w[+mC]=r4-GAL4}3* (RRID:BDSC_33832). Fat body-specific dERR rescue experiments were conducted using a previously described strain harboring a *UAS-dERR* transgene (Tennessen et al., 2011). RNAi experiments were conducted using y1 v1; P{y+t7.7v[+t1.8]=TRiP.HMC03087}attP2 (RRID:BDSC_50686), which was generated by the *Drosophila* Transgenic RNAi Project (Zirin et al., 2020). The strain *w^1118^; PBac{y[+mDint2] w[+mC]=ERR-GFP.FSTF}VK00037* (RRID:BDSC_38638) was used to examine dERR expression in the larval fat body. This strain harbors an integrated bacterial artificial chromosome that contains the *dERR* genomic locus with a GFP-StrepII-FLAG TAG inserted at the dERR C-terminus (Venken et al., 2009). dHNF4-LBD activation experiments were conducted using the previously described *hs-GAL4-dHNF4* transgene (Palanker et al., 2006). Flybase was used as a reference throughout the study (Öztürk-Çolak et al., 2024).

### Immunofluorescence

Larvae were dissected in 1x phosphate buffered saline (PBS) and fixed for 30 minutes in 4% paraformaldehyde in 1x PBS at room temperature. Fixed tissues were then washed 3 times for 5 minutes each with PT (PBS with 0.1% Triton X-100) and blocked for 30 minutes in NGS-PT (PT with 5% normal goat serum (NGS)) at room temperature. Blocked tissues were incubated overnight at 4 °C with Mouse anti-FLAG (# MA1-91878 Thermo Fisher) diluted 1:1000 in NGS-PT. Tissues were subsequently washed 3 times for 5 minutes each with PT and then incubated overnight with Alexa Fluor 488 Goat anti-Mouse secondary antibody (#A11029 Thermo Fisher) diluted 1:1000 in PT at 4 °C. Tissues were then washed 3 times for 5 minutes each in PT and mounted in Vectashield with DAPI (Vector Laboratories; H-1200-314 10).

### Fat Body Staining

Nile red staining was conducted on L2 fat bodies as previously described (Grönke et al., 2005). Briefly, dissected tissues were fixed with 4% paraformaldehyde in PBS for 30 min and washed with PBST three times for 5 minutes each, and then incubated for 1 hr at room temperature in a 1:1000 dilution of Nile Red stock (10mg/mL in acetone) in 1X PBS. Tissues were washed three times for 5 minutes each in PBST, mounted in Vectashield (Vector Laboratories), and imaged on a Leica SP8 Confocal at 568nm.

Solvent Black 3 staining was conducted as described. Briefly, fat bodies were fixed with 4% formaldehyde, rinsed twice with PBS, once with 50% ethanol, and then stained for 2 minutes at room temperature using filtered 0.5% Solvent Black 3 (CAS Number 4197-25-5; Sigma 199664) dissolved in 75% ethanol. Samples were sequentially rinsed with 50% ethanol, 25% ethanol, and PBS. Stained tissues were mounted on a microscope slide with vector shield with DAPI (Vector Laboratories; H-1200-10).

### TAG measurements

TAG was quantified from larval extracts as described (Tennessen et al., 2014). Briefly, twenty-five mid-L2 larvae of the appropriate genotypes were then collected in 1.5 mL microfuge tubes, washed three times with phosphate-buffered saline pH 7.0 (PBS), homogenized in 100 µl of cold PBS + 0.05% Tween 20 (PBST), and heat-treated for 10 min at 70°C. The resulting homogenate was assayed for triglyceride and soluble protein levels as previously described (Tennessen et al., 2014).

### HNF4 Ligand Binding Domain Assays

Analysis was conducted using either *w^1118^; hs-GAL4-dHNF4 UAS-nlacZ; dERR^1^* mutants, *w^1118^; hs-GAL4-dHNF4 UAS-nlacZ; dERR^1/+^* heterozygous controls, or *w^1118^; UAS-nlacZ* negative control larvae. Tissues from mid-L2 larvae were dissected and fixed as described (Palanker et al., 2006). Briefly, molasses-yeast plates containing mid-L2 larvae were heat-treated for 30 minutes at 37°C and then allowed to recover for 2 hours at 25°C. Larvae were dissected in 1x PBS, fixed for 20 minutes in 4% PFA in 1x PBS at room temperature, and stained in 0.2% Xgal staining solution for 2 hrs at 37°C. Following the staining period, tissues were washed 3 times for 5 min each in 1x PBS, mounted in a solution of 50% glycerol, and imaged immediately following mounting.

### Lipidomic Analysis

#### Chemicals

LC-MS-grade solvents and mobile phase modifiers were obtained from Honeywell Burdick & Jackson, Morristown, NJ (acetonitrile, isopropanol, formic acid), Fisher Scientific, Waltham, MA (methyl *tert*-butyl ether) and Sigma–Aldrich/Fluka, St. Louis, MO (ammonium formate, ammonium acetate). Lipid standards were obtained from Avanti Polar Lipids, Alabaster, AL (EquiSPLASH LIPIDOMIX (330731)) and Cayman Chemical, Ann Arbor, MI, (Palmitic acid d-31 (16497)).

#### Sample Preparation

Extraction of lipids was carried out using a biphasic solvent system of cold methanol, methyl *tert*-butyl ether (MTBE), PBS, and water as previously described with some modifications (Matyash et al., 2008). In a randomized sequence, to each sample was added 225 µL MeOH with internal standards and 188 µL PBS. Samples were homogenized for 30 seconds, transferred to 13x100 mm screw-capped glass test tubes containing 750 µL MTBE, and then incubated on ice with occasional vortexing for 1 hr. Following incubation, samples were centrifuged at 15,000 x g for 10 minutes at 4 °C. The organic (upper) layer was collected, and the aqueous (lower) layer was re-extracted with 1 mL of 10:3:2.5 (*v/v/v*) MTBE/MeOH/dd-H2O, briefly vortexed, incubated at RT, and centrifuged at 15,000 x g for 10 minutes at 4 °C. Upper phases were combined and evaporated to dryness under nitrogen. Lipid extracts were reconstituted in 800 µL of 4:1:1 (v/v/v) IPA/ACN/water and transferred to LC-MS vials for analysis. Concurrently, a process blank sample was prepared and pooled quality control (QC) samples were prepared by taking equal volumes from each sample after final resuspension.

#### Mass Spectrometry Analysis of Samples

Lipid extracts were separated on an Acquity UPLC CSH C18 column (2.1 x 100 mm; 1.7 µm) coupled to an Acquity UPLC CSH C18 VanGuard precolumn (5 × 2.1 mm; 1.7 µm) (Waters, Milford, MA) maintained at 65 °C connected to an Agilent HiP 1290 Sampler, Agilent 1290 Infinity pump, and Agilent 6545 Accurate Mass Q-TOF dual AJS-ESI mass spectrometer (Agilent Technologies, Santa Clara, CA). Samples were analyzed in a randomized order in both positive and negative ionization modes in separate experiments acquiring with the scan range m/z 100 – 1700. For positive mode, the source gas temperature was set to 225 °C, with a drying gas flow of 11 L/min, nebulizer pressure of 40 psig, sheath gas temp of 350 °C and sheath gas flow of 11 L/min. VCap voltage is set at 3500 V, nozzle voltage 500V, fragmentor at 110 V, skimmer at 85 V and octopole RF peak at 750 V. For negative mode, the source gas temperature was set to 300 °C, with a drying gas flow of 11 L/min, a nebulizer pressure of 30 psig, sheath gas temp of 350 °C and sheath gas flow 11 L/min. VCap voltage was set at 3500 V, nozzle voltage 75 V, fragmentor at 175 V, skimmer at 75 V and octopole RF peak at 750 V. Mobile phase A consisted of ACN:H_2_O (60:40, *v/v*) in 10 mM ammonium formate and 0.1% formic acid, and mobile phase B consisted of IPA:ACN:H_2_O (90:9:1, *v/v/v*) in 10 mM ammonium formate and 0.1% formic acid. For negative mode analysis the modifiers were changed to 10 mM ammonium acetate. The chromatography gradient for both positive and negative modes started at 15% mobile phase B then increased to 30% B over 2.4 min, it then increased to 48% B from 2.4 – 3.0 min, then increased to 82% B from 3 – 13.2 min, then increased to 99% B from 13.2 – 13.8 min where it was held until 16.7 min, then returned to the initial conditions and equilibriated for 5 min. Flow was 0.4 mL/min throughout, with injection volumes of 1 µL for positive and 10 µL negative mode. Tandem mass spectrometry was conducted using iterative exclusion, with the same LC gradient at collision energies of 20 V and 27.5 V in positive and negative modes, respectively.

#### Analysis of Mass Spectrometry Data

For data processing, Agilent MassHunter (MH) Workstation and software packages MH Qualitiative and MH Quantitative were used. The pooled QC (n=8) and process blank (n=4) were injected throughout the sample queue to ensure the reliability of acquired lipidomics data. For lipid annotation, accurate mass and MS/MS matching was used with the Agilent Lipid Annotator library and LipidMatch (Koelmel et al., 2017). Results from the positive and negative ionization modes from Lipid Annotator were merged based on the class of lipid identified. Data exported from MH Quantitative was evaluated using Excel where initial lipid targets are parsed based on the following criteria. Only lipids with relative standard deviations (RSD) less than 30% in QC samples are used for data analysis. Additionally, only lipids with background AUC counts in process blanks that are less than 30% of QC are used for data analysis. The parsed excel data tables are normalized based on the ratio to class-specific internal standards, then to tissue mass prior to statistical analysis.

#### Statistical Analysis and Data Visualization

Multivariate analysis was performed using MetaboAnalyst (Pang et al., 2024). Statistical models were created for the normalized data after logarithmic transformation (base 10) and Pareto scaling. Initial pass for the volcano plot used a fold change (FC) cut off of 1.5, adjusted p-value cut off: 0.05, and multiple testing correction: false discovery rate.

### Sample preparation and RNA extraction for RNA-Seq

*dERR^1/+^ and dERR^1/2^* L2 larvae (66-68 hours) were raised at 25°C on molasses plates covered with yeast. Larval stage was confirmed by spiracle morphology prior to dissection. Larvae were washed and dissected in 1x PBS in groups of 5 to ensure speed of dissection and preservation of RNA quality. A total of 15 larvae were dissected to make three sets of technical replicates per genotype. No. 5 Biology Grade Dumont INOX dissecting forceps were used to decapitate larvae and dissect the fat body from surrounding tissue. Complete RNA was extracted using RNeasy Plus Micro Kit (74034, Qiagen) following manufacturer’s instructions.The fat body was quickly moved to a 1.7ml Eppendorf tube with a small volume (50 ml) of the RLT Plus Buffer with beta-mercaptoethanol added. The tissue was homogenized in the solution with a sterile pestle while keeping the tube on ice. The homogenized samples were flash frozen in liquid nitrogen and stored at -80°C until further processing. After removal of samples from the - 80°C, an additional 300uL of RLT Plus was added to the sample just before it completely thawed. Then 350ul of 70% ethanol was added and the combined 700ul pipetted forcefully to mix. Immediately thereafter, the sample was transferred to a RNeasy MinElute spin column before proceeding with the rest of the manufacturer protocol (starting at RNAeasy kit step 5).

### RNA-seq Data Processing

RNA-seq processing scripts are included in Figure S2 and all commands are run with default settings unless otherwise stated. FastQC (0.12.1) was used to assess read quality prior to further processing (Andrews, 2010). The D. melanogaster BDGP6.46 hard-masked assembly (Dainat, 2024) and v110 annotation (Pertea and Pertea, 2020) were downloaded from ENSEMBL, and malformed entries in the annotation file were corrected using AGAT (1.2.0) ‘agat convert sp gxf2gxf.pl’ (Dainat, 2024). gffread (0.12.7) was used to create a transcript sequence file (Pertea and Pertea, 2020). The genome assembly was concatenated, followed by decoy-aware indexing with Salmon (1.10.2) (Patro et al., 2017). Salmon quantification was then performed with non-default settings: ‘--seqBias --gcBias --posBias --softclip softclipOverhangs -- numGibbsSamples 100’.

### RNA-seq Downstream Analysis

The downstream RNA-seq analysis scripts are included in Figure S2 and all analyses were performed with R (4.3.2). Salmon transcript abundance estimates were aggregated to gene level and imported into R using tximport (1.30.0) (Soneson et al., 2015). For quality control, PCAtools (2.14.0, Blighe and Lun, 2022) was utilized for PCA analysis, using DESeq2 (1.42.0, Love et al., 2014) ‘rlog()’-normalized counts with the design ‘∼ condition + source + condition:source’ and blind=FALSE, where ‘condition’ is the ERR mutation status and ‘source’ the tissue source type (i.e. whole body or fat body). For differential expression, genes were considered to have a significant interaction term, i.e. their log2FC is significantly different depending on whether the tissue was from the whole body or fat body, if the FDR < 0.05. Separate contrasts were performed with a simplified design (‘∼ condition_source’ where ‘condition_source’ is the concatenated ‘condition’ and ‘source’ factor levels) to compare whole body or fat body ERR mutant versus controls, and apeglm (1.24.0, Zhu et al., 2019) was used for log fold change shrinkage as well as testing against |log2FC| > 1, with significant genes having an s-value < 0.005. An additional analysis was performed where only genes that were both differentially expressed (s-value < 0.005) and with a significant interaction term (FDR < 0.05) were considered. Volcano and barplots were generated with ggplot2 (3.4.4), euler plots with eulerr (7.0.0,Wilkinson, 2012), and heatmaps with ComplexHeatmap (2.18.0, Gu et al., 2016). The ‘fora()’ function from fgsea (1.28.0, Korotkevich et al., 2021) was used to perform hypergeometric enrichment analysis using the following gene sets pulled from Molecular Signatures Database (MSigDB, Subramanian et al., 2005) via the msigdbr (7.5.1, Dolgalev, 2022) library: hallmark gene sets, chemical and genetic perturbations, reactome pathways, gene ontology molecular function and biological process, and human phenotype ontology. Enrichment plots were generated using ggplot2.

### Gene Ontology Analysis using PANGEA

RNAseq data were analyzed using PANGEA (Hu et al., 2023). Genes that were significantly down- or up-regulated were analyzed for Gene Ontology Enrichment using the SLIM2 GO BP and FlyBase signaling pathway (experimental evidence) sets (Consortium et al., 2023).

## Data Availability

Processed RNA-seq data is presented in Table S4 and S5. Original data is available in NCBI Gene Expression Omnibus (GEO; GSE 273774).

## ACKNOWLEDGEMENTS

We thank the Bloomington *Drosophila* Stock Center (NIH P40OD018537) for providing fly stocks, the *Drosophila* Genomics Resource Center (NIH 2P40OD010949) for genomic reagents, Flybase (NIH 5U41HG000739), the Indiana University Light Microscopy Imaging Center, and the Indiana University Center for Genomics and Bioinformatics. Metabolomics analysis was performed at the Metabolomics Core Facility at the University of Utah. Mass spectrometry equipment was obtained through NCRR Shared Instrumentation Grant 1S10OD016232-01, 1S10OD018210-01A1 and 1S10OD021505-01. Authors thank Dan Cuthbertson of Agilent Technologies for assistance in implementing iterative exclusion in the tandem mass spectrometry experiments. J.M.T. is supported by the National Institute of General Medical Sciences of the National Institutes of Health under a R35 Maximizing Investigators’ Research Award (MIRA; 1R35GM119557).

## SUPPLEMENTAL FIGURE LEGENDS

**Figure S1. A comparison of the lipidomic data from *ERR^1/2^*mutants and *ERR^1/+^* control samples using principal component (PC) analysis.** Targeted metabolomics data from Table S1 was analyzed using principal component analysis. Analysis was conducted using Metaboanalyst 5.0.

**Figure S2. R scripts used to conduct the RNA-seq processing and RNA-seq analysis.**

## Supplemental Tables

**Table S1.** Lipidomic analysis of *dERR^1/2^* mutants compared with *dERR^1/+^*heterozygous controls. Samples were collected 60-hours after egg laying (mid L2). Each sample consisted of 20 larvae. Concentrations listed as pmol/mg.

**Table S2.** A list of lipids that exhibit significant changes in *dERR^1/2^* mutants compared with ERR^1/+^ heterozygous controls (absolute FC>2, adjusted *P*<0.1).

**Table S3.** A list of lipids that exhibit significant changes in *dERR^1/2^* mutants compared with *dERR^1/+^* heterozygous controls (absolute FC>1.5, adjusted *P*<0.05).

**Table S4.** RNAseq analysis comparing gene expression in the whole bodies of *dERR^1/2^* mutants with *dERR^1/+^* heterozygous controls. Larvae were collected 60-hours after egg laying (mid L2).

**Table S5.** RNAseq analysis comparing gene expression in fat bodies isolated from *dERR^1/2^*mutants with *dERR^1/+^*heterozygous controls. Larvae were collected 60-hours after egg laying (mid L2).

**Table S6.** Genes that are significantly down-regulated in either Table S4 (*dERR* mutant RNA-seq; whole body) or Table S5 (*dERR* mutant RNA-seq; fat body) are listed according to whether they are down-regulated in only whole animal samples, only fat body samples, or in both sample sets. All genes listed displayed a log2 fold change<-1 and a sval<0.005.

**Table S7.** Genes that are significantly up-regulated in either Table S4 (*dERR* mutant RNA-seq; whole body) or Table S5 (*dERR* mutant RNA-seq; fat body) are listed according to whether they are down-regulated in only whole animal samples, only fat body samples, or in both sample sets. All genes listed displayed a log2 fold change>1 and a sval<0.005.

**Table S8.** PANGEA analysis of significantly down-regulated genes (log2 fold change≤-1 and a sval<0.005) in the fat body of *dERR^1/2^* mutants as compared with *dERR^1/+^* heterozygous controls (see Table S5). PANGEA default settings (FlyBase signaling pathway; SLIM2 GO BP) were used to analyze for enrichment. Only Gene Set IDs with a *P-*value of less than 0.1 are included in the table.

**Table S9.** PANGEA analysis of significantly up-regulated genes (log2 fold change≤-1 and a sval<0.005) in the fat body of *dERR^1/2^* mutants as compared with *dERR^1/+^* heterozygous controls (see Table S5). PANGEA default settings (FlyBase signaling pathway; SLIM2 GO BP) were used to analyze for enrichment. Only Gene Set IDs with a *P-*value of less than 0.1 are included in the table.

**Table S10.** Expression of genes encoding beta-oxidation enzymes in *dERR* mutants.

## Literature Cited

Aguila, J. R., Suszko, J., Gibbs, A. G. & Hoshizaki, D. K. 2007. The role of larval fat cells in adult Drosophila melanogaster. J Exp Biol, 210, 956–63.

Andrews, S. 2010. FastQC: a quality control tool for high throughput sequence data. [Online]. Available: http://www.bioinformatics.babraham.ac.uk/projects/fastqc [Accessed].

Baker, E. R. 1985. Body weight and the initiation of puberty. Clin Obstet Gynecol, 28, 573–9.

Barrea, L., Vetrani, C., Verde, L., FRIAS-Toral, E., GARCIA-Velasquez, E., Ranasinghe, P., Mendez, V., Jayawardena, R., Savastano, S., Colao, A. & Muscogiuri, G. 2022. Gestational obesity: An unconventional endocrine disruptor for the fetus. Biochem Pharmacol, 198, 114974.

Barton, L. J., Sanny, J., PACKARD Dawson, E., Nouzova, M., Noriega, F. G., Stadtfeld, M. & Lehmann, R. 2024. Juvenile hormones direct primordial germ cell migration to the embryonic gonad. Current Biology, 34, 505–518.e6.

Beebe, K., Robins, M. M., Hernandez, E. J., Lam, G., Horner, M. A. & Thummel, C. S. 2020. Drosophila estrogen-related receptor directs a transcriptional switch that supports adult glycolysis and lipogenesis. Genes Dev, 34, 701–714.

Bellés, X., Martín, D. & Piulachs, M. D. 2005. The mevalonate pathway and the synthesis of juvenile hormone in insects. Annu Rev Entomol, 50, 181–99.

Bi, J., Xiang, Y., Chen, H., Liu, Z., Grönke, S., Kühnlein, R. P. & Huang, X. 2012. Opposite and redundant roles of the two Drosophila perilipins in lipid mobilization. Journal of Cell Science, 125, 3568–3577.

Blighe, K. & Lun, A. 2022. PCAtools: Everything Principal Components Analysis. R package version 2.8.0 [Online]. Available: https://github.com/kevinblighe/PCAtools [Accessed].

Carrier, J. C., Deblois, G., Champigny, C., Levy, E. & Giguère, V. 2004. Estrogen-related receptor alpha (ERRalpha) is a transcriptional regulator of apolipoprotein A-IV and controls lipid handling in the intestine. J Biol Chem, 279, 52052–8.

Carvalho, M., Sampaio, J. L., Palm, W., Brankatschk, M., Eaton, S. & Shevchenko, A. 2012. Effects of diet and development on the Drosophila lipidome. Mol Syst Biol, 8, 600.

Church, R. B. & Robertson, F. W. 1966. A biochemical study of the growth of *Drosophila melanogaster*. J. Exp. Zool., 162, 337–351.

Consortium, T. G. O., Aleksander, S. A., Balhoff, J., Carbon, S., Cherry, J. M., Drabkin, H. J., Ebert, D., Feuermann, M., Gaudet, P., Harris, N. L., Hill, D. P., Lee, R., Mi, H., Moxon, S., Mungall, C. J., Muruganugan, A., Mushayahama, T., Sternberg, P. W., Thomas, P. D., VAN Auken, K., Ramsey, J., Siegele, D. A., Chisholm, R. L., Fey, P., Aspromonte, M. C., Nugnes, M. V., Quaglia, F., Tosatto, S., Giglio, M., Nadendla, S., Antonazzo, G., Attrill, H., DOS Santos, G., Marygold, S., Strelets, V., Tabone, C. J., Thurmond, J., Zhou, P., Ahmed, S. H., Asanitthong, P., LUNA Buitrago, D., Erdol, M. N., Gage, M. C., ALI Kadhum, M., Li, K. Y. C., Long, M., Michalak, A., Pesala, A., Pritazahra, A., Saverimuttu, S. C. C., Su, R., Thurlow, K. E., Lovering, R. C., Logie, C., Oliferenko, S., Blake, J., Christie, K., Corbani, L., Dolan, M. E., Drabkin, H. J., Hill, D. P., Ni, L., Sitnikov, D., Smith, C., Cuzick, A., Seager, J., Cooper, L., Elser, J., Jaiswal, P., Gupta, P., Jaiswal, P., Naithani, S., LERA-Ramirez, M., Rutherford, K., Wood, V., DE Pons, J. L., Dwinell, M. R., Hayman, G. T., Kaldunski, M. L., Kwitek, A. E., Laulederkind, S. J. F., Tutaj, M. A., Vedi, M., Wang, S.-J., D’eustachio, P., Aimo, L., Axelsen, K., Bridge, A., HYKA-Nouspikel, N., Morgat, A., Aleksander, S. A., Cherry, J. M., Engel, S. R., Karra, K., Miyasato, S. R., Nash, R. S., Skrzypek, M. S., Weng, S., Wong, E. D., Bakker, E., et al. 2023. The Gene Ontology knowledgebase in 2023. Genetics, 224.

Dainat, J. 2024. NBISweden/AGAT: AGAT-v1.4.0. v1.4.0 ed. Zenodo.

Danielsen, E. T., Moeller, M. E. & Rewitz, K. F. 2013. Nutrient signaling and developmental timing of maturation. Curr Top Dev Biol, 105, 37–67.

Deshpande, G., Godishala, A. & Schedl, P. 2009. Ggamma1, a downstream target for the hmgcr-isoprenoid biosynthetic pathway, is required for releasing the Hedgehog ligand and directing germ cell migration. PLoS Genet, 5, e1000333.

Deshpande, G. & Schedl, P. 2005. *HMGCoA reductase* Potentiates *hedgehog* Signaling in *Drosophila melanogaster*. Developmental Cell, 9, 629-638.

Deshpande, G., Zhou, K., Wan, J. Y., Friedrich, J., Jourjine, N., Smith, D. & Schedl, P. 2013. The hedgehog pathway gene shifted functions together with the hmgcr-dependent isoprenoid biosynthetic pathway to orchestrate germ cell migration. PLoS Genet, 9, e1003720.

Dolgalev, I. 2022. msigdbr: MSigDB Gene Sets for Multiple Organisms in a Tidy Data Format.. 7.5.1 ed.

Gáliková, M. & Klepsatel, P. 2023. Endocrine control of glycogen and triacylglycerol breakdown in the fly model. Semin Cell Dev Biol, 138, 104–116.

Garfinkel, A. M., Ilker, E., Miyazawa, H., Schmeisser, K. & Tennessen, J. M. 2024. Historic obstacles and emerging opportunities in the field of developmental metabolism - lessons from Heidelberg. Development, 151.

Garrido, D., Rubin, T., Poidevin, M., Maroni, B., LE Rouzic, A., Parvy, J. P. & Montagne, J. 2015. Fatty acid synthase cooperates with glyoxalase 1 to protect against sugar toxicity. PLoS Genet, 11, e1004995.

Geng, D. Q., Wang, X. L., Lyu, X. Y., Raikhel, A. S. & Zou, Z. 2024. Ecdysone-controlled nuclear receptor ERR regulates metabolic homeostasis in the disease vector mosquito Aedes aegypti. PLoS Genet, 20, e1011196.

Grant, J., Saldanha, J. W. & Gould, A. P. 2010. A Drosophila model for primary coenzyme Q deficiency and dietary rescue in the developing nervous system. Dis Model Mech, 3, 799–806.

Grönke, S., Mildner, A., Fellert, S., Tennagels, N., Petry, S., Müller, G., Jäckle, H. & Kühnlein, R. P. 2005. Brummer lipase is an evolutionary conserved fat storage regulator in Drosophila. Cell Metab, 1, 323–30.

Gu, Z., Eils, R. & Schlesner, M. 2016. Complex heatmaps reveal patterns and correlations in multidimensional genomic data. Bioinformatics, 32, 2847–9.

Heier, C., Klishch, S., Stilbytska, O., Semaniuk, U. & Lushchak, O. 2021. The Drosophila model to interrogate triacylglycerol biology. Biochim Biophys Acta Mol Cell Biol Lipids, 1866, 158924.

Ho, S.-Y., Thorpe, J. L., Deng, Y., Santana, E., Derose, R. A. & Farber, S. A. 2004. Lipid Metabolism in Zebrafish. Methods in Cell Biology. Academic Press.

Hofbauer, H. F., Heier, C., SEN Saji, A. K. & Kühnlein, R. P. 2021. Lipidome remodeling in aging normal and genetically obese Drosophila males. Insect Biochem Mol Biol, 133, 103498.

Hu, Y., Comjean, A., Attrill, H., Antonazzo, G., Thurmond, J., Chen, W., Li, F., Chao, T., Mohr, S. E., Brown, N. H. & Perrimon, N. 2023. PANGEA: a new gene set enrichment tool for Drosophila and common research organisms. Nucleic Acids Research, 51, W419–W426.

Huang, L., Hou, J. W., Fan, H. Y., Tsai, M. C., Yang, C., Hsu, J. B. & Chen, Y. C. 2023. Critical body fat percentage required for puberty onset: the Taiwan Pubertal Longitudinal Study. J Endocrinol Invest, 46, 1177–1185.

Koelmel, J. P., Kroeger, N. M., Ulmer, C. Z., Bowden, J. A., Patterson, R. E., Cochran, J. A., Beecher, C. W. W., Garrett, T. J. & Yost, R. A. 2017. LipidMatch: an automated workflow for rule-based lipid identification using untargeted high-resolution tandem mass spectrometry data. BMC Bioinformatics, 18, 331.

Korotkevich, G., Sukhov, V., Budin, N., Shpak, B., Artyomov, M. N. & Sergushichev, A. 2021. Fast gene set enrichment analysis. bioRxiv, 060012.

Lehmann, M. 2018. Endocrine and physiological regulation of neutral fat storage in Drosophila. Mol Cell Endocrinol, 461, 165–177.

Li, H. & Tennessen, J. M. 2017. Methods for studying the metabolic basis of Drosophila development. Wiley Interdiscip Rev Dev Biol, 6.

Li, S., Yu, X. & Feng, Q. 2019. Fat Body Biology in the Last Decade. Annu Rev Entomol, 64, 315–333.

Li, Y., Ma, T., Ma, Y., Gao, D., Chen, L., Chen, M., Liu, J., Dong, B., Dong, Y. & Ma, J. 2022. Adiposity Status, Trajectories, and Earlier Puberty Onset: Results From a Longitudinal Cohort Study. J Clin Endocrinol Metab, 107, 2462–2472.

Long, W., Wu, J., Shen, G., Zhang, H., Liu, H., Xu, Y., Gu, J., Jia, L., Lin, Y. & Xia, Q. 2020. Estrogen-related receptor participates in regulating glycolysis and influences embryonic development in silkworm Bombyx mori. Insect Mol Biol, 29, 160–169.

Love, M. I., Huber, W. & Anders, S. 2014. Moderated estimation of fold change and dispersion for RNA-seq data with DESeq2. Genome Biol, 15, 550.

Luo, J., Sladek, R., Carrier, J., Bader, J. A., Richard, D. & Giguère, V. 2003. Reduced fat mass in mice lacking orphan nuclear receptor estrogen-related receptor alpha. Mol Cell Biol, 23, 7947–56.

Marcus, C., Danielsson, P. & Hagman, E. 2022. Pediatric obesity-Long-term consequences and effect of weight loss. J Intern Med, 292, 870–891.

Matyash, V., Liebisch, G., Kurzchalia, T. V., Shevchenko, A. & Schwudke, D. 2008. Lipid extraction by methyl-tert-butyl ether for high-throughput lipidomics. J Lipid Res, 49, 1137–46.

Menendez, A., Wanczyk, H., Walker, J., Zhou, B., Santos, M. & Finck, C. 2022. Obesity and Adipose Tissue Dysfunction: From Pediatrics to Adults. Genes (Basel), 13.

Miyazawa, H. & Aulehla, A. 2018. Revisiting the role of metabolism during development. Development, 145.

Musselman, L. P., Fink, J. L., Ramachandran, P. V., Patterson, B. W., Okunade, A. L., Maier, E., Brent, M. R., Turk, J. & Baranski, T. J. 2013. Role of fat body lipogenesis in protection against the effects of caloric overload in Drosophila. J Biol Chem, 288, 8028–8042.

Musselman, L. P. & Kühnlein, R. P. 2018. Drosophila as a model to study obesity and metabolic disease. Journal of Experimental Biology, 221.

ÖZTÜRK-Çolak, A., Marygold, S. J., Antonazzo, G., Attrill, H., GOUTTE-Gattat, D., Jenkins, V. K., Matthews, B. B., Millburn, G., DOS Santos, G. & Tabone, C. J. 2024. FlyBase: updates to the Drosophila genes and genomes database. Genetics, 227.

PALACIOS-Marin, I., Serra, D., JIMÉNEZ-Chillarón, J. C., Herrero, L. & Todorčević, M. 2023. Childhood obesity: Implications on adipose tissue dynamics and metabolic health. Obesity Reviews, 24, e13627.

Palanker, L., Necakov, A. S., Sampson, H. M., Ni, R., Hu, C., Thummel, C. S. & Krause, H. M. 2006. Dynamic regulation of Drosophila nuclear receptor activity in vivo. Development, 133, 3549–62.

Palanker, L., Tennessen, J. M., Lam, G. & Thummel, C. S. 2009. Drosophila HNF4 regulates lipid mobilization and beta-oxidation. Cell Metab, 9, 228–39.

Palm, W. & Rodenfels, J. 2020. Understanding the role of lipids and lipoproteins in development. Development, 147.

Pang, Z., Lu, Y., Zhou, G., Hui, F., Xu, L., Viau, C., Spigelman, ALIYA F., Macdonald, PATRICK E., Wishart, DAVID S., Li, S. & Xia, J. 2024. MetaboAnalyst 6.0: towards a unified platform for metabolomics data processing, analysis and interpretation. Nucleic Acids Research, 52, W398–W406.

Patro, R., Duggal, G., Love, M. I., Irizarry, R. A. & Kingsford, C. 2017. Salmon provides fast and bias-aware quantification of transcript expression. Nat Methods, 14, 417–419.

Pertea, G. & Pertea, M. 2020. GFF Utilities: GffRead and GffCompare. F1000Res, 9.

Rodenfels, J., Lavrynenko, O., Ayciriex, S., Sampaio, J. L., Carvalho, M., Shevchenko, A. & Eaton, S. 2014. Production of systemically circulating Hedgehog by the intestine couples nutrition to growth and development. Genes Dev, 28, 2636–51.

Santos, A. C. & Lehmann, R. 2004. Isoprenoids Control Germ Cell Migration Downstream of HMGCoA Reductase. Developmental Cell, 6, 283–293.

Scheidl, T. B., Brightwell, A. L., Easson, S. H. & Thompson, J. A. 2023. Maternal obesity and programming of metabolic syndrome in the offspring: searching for mechanisms in the adipocyte progenitor pool. BMC Med, 21, 50.

Scholtes, C., Dufour, C. R., Pleynet, E., Kamyabiazar, S., Hutton, P., Baby, R., Guluzian, C. & Giguère, V. 2024. Identification of a chromatin-bound ERRα interactome network in mouse liver. Mol Metab, 83, 101925.

Simcox, J., Geoghegan, G., Maschek, J. A., Bensard, C. L., Pasquali, M., Miao, R., Lee, S., Jiang, L., Huck, I., Kershaw, E. E., Donato, A. J., Apte, U., Longo, N., Rutter, J., Schreiber, R., Zechner, R., Cox, J. & Villanueva, C. J. 2017. Global Analysis of Plasma Lipids Identifies Liver-Derived Acylcarnitines as a Fuel Source for Brown Fat Thermogenesis. Cell Metab, 26, 509–522.e6.

Skowronek, P., Wójcik, Ł. & Strachecka, A. 2021. Fat Body-Multifunctional Insect Tissue. Insects, 12.

Soneson, C., Love, M. I. & Robinson, M. D. 2015. Differential analyses for RNA-seq: transcript-level estimates improve gene-level inferences. F1000Res, 4, 1521.

Srinivasan, S. 2015. Regulation of body fat in Caenorhabditis elegans. Annu Rev Physiol, 77, 161–78.

Subramanian, A., Tamayo, P., Mootha, V. K., Mukherjee, S., Ebert, B. L., Gillette, M. A., Paulovich, A., Pomeroy, S. L., Golub, T. R., Lander, E. S. & Mesirov, J. P. 2005. Gene set enrichment analysis: a knowledge-based approach for interpreting genome-wide expression profiles. Proc Natl Acad Sci U S A, 102, 15545–50.

Suh, J. M., Gao, X., Mckay, J., Mckay, R., Salo, Z. & Graff, J. M. 2006. Hedgehog signaling plays a conserved role in inhibiting fat formation. Cell Metab, 3, 25–34.

Tennessen, J. M., Baker, K. D., Lam, G., Evans, J. & Thummel, C. S. 2011. The Drosophila estrogen-related receptor directs a metabolic switch that supports developmental growth. Cell Metab, 13, 139–48.

Tennessen, J. M., Barry, W. E., Cox, J. & Thummel, C. S. 2014. Methods for studying metabolism in Drosophila. Methods, 68, 105–15.

Tennessen, J. M. & Thummel, C. S. 2011. Coordinating growth and maturation - insights from Drosophila. Curr Biol, 21, R750–7.

Texada, M. J., Koyama, T. & Rewitz, K. 2020. Regulation of Body Size and Growth Control. Genetics, 216, 269–313.

Venken, K. J., Carlson, J. W., Schulze, K. L., Pan, H., He, Y., Spokony, R., Wan, K. H., Koriabine, M., DE Jong, P. J., White, K. P., Bellen, H. J. & Hoskins, R. A. 2009. Versatile P[acman] BAC libraries for transgenesis studies in Drosophila melanogaster. Nat Methods, 6, 431–4.

Watts, J. L. & Ristow, M. 2017. Lipid and Carbohydrate Metabolism in Caenorhabditis elegans. Genetics, 207, 413–446.

Wilkinson, L. 2012. Exact and approximate area-proportional circular Venn and Euler diagrams. IEEE Trans Vis Comput Graph, 18, 321–31.

Yi, P., Han, Z., Li, X. & Olson, E. N. 2006. The mevalonate pathway controls heart formation in Drosophila by isoprenylation of Ggamma1. Science, 313, 1301–3.

Zhang, J., Liu, Y., Jiang, K. & Jia, J. 2020. Hedgehog signaling promotes lipolysis in adipose tissue through directly regulating Bmm/ATGL lipase. Dev Biol, 457, 128–139.

Zhu, A., Ibrahim, J. G. & Love, M. I. 2019. Heavy-tailed prior distributions for sequence count data: removing the noise and preserving large differences. Bioinformatics, 35, 2084–2092.

Zhu, H. & Han, M. 2014. Exploring developmental and physiological functions of fatty acid and lipid variants through worm and fly genetics. Annu Rev Genet, 48, 119–48.

Zirin, J., Hu, Y., Liu, L., YANG-Zhou, D., Colbeth, R., Yan, D., EWEN-Campen, B., Tao, R., Vogt, E., Vannest, S., Cavers, C., Villalta, C., Comjean, A., Sun, J., Wang, X., Jia, Y., Zhu, R., Peng, P., Yu, J., Shen, D., Qiu, Y., Ayisi, L., Ragoowansi, H., Fenton, E., Efrem, S., Parks, A., Saito, K., Kondo, S., Perkins, L., Mohr, S. E., Ni, J. & Perrimon, N. 2020. Large-Scale Transgenic Drosophila Resource Collections for Loss- and Gain-of-Function Studies. Genetics, 214, 755–767.

