## Supplemental Figure 1 for "The *Drosophila* Estrogen-Related Receptor promotes triglyceride storage within the larval fat body"

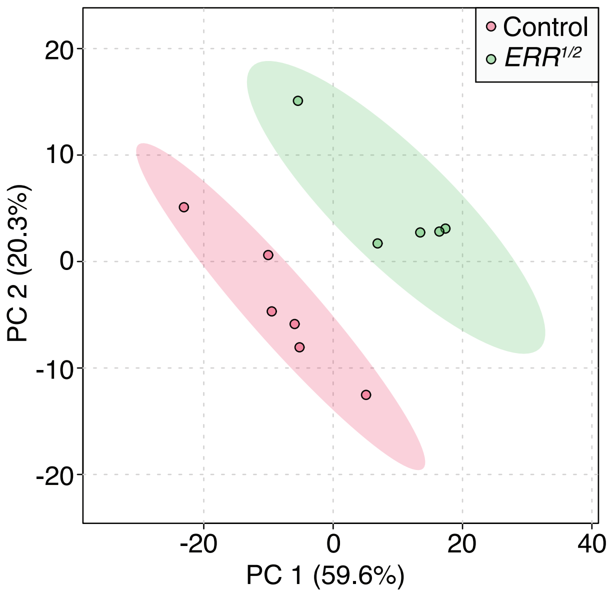


**Figure S1.** A comparison of the lipidomic data from ERR1/2 mutants and ERR1/+ control samples using principal component (PC) analysis. Targeted metabolomics data from Table S1 was analyzed using principal component analysis. Analysis was conducted using Metaboanalyst 5.0.
