## Supplemental Figure 2 for "The *Drosophila* Estrogen-Related Receptor promotes triglyceride storage within the larval fat body"

### Tennessen RNA-seq

Bob Policastro

2023-12-29

Code

#### Libraries

Load the required libraries.

Hide

```
library("tidyverse")
#> ── Attaching core tidyverse packages ──────────────────────────────── tidyverse 2.0.0 ──
#> ✔ dplyr     1.1.4     ✔ readr     2.1.5
#> ✔ forcats   1.0.0     ✔ stringr   1.5.1
#> ✔ ggplot2   3.4.4     ✔ tibble    3.2.1
#> ✔ lubridate 1.9.3     ✔ tidyr     1.3.1
#> ✔ purrr     1.0.2
#> ── Conflicts ────────────────────────────────────────── tidyverse_conflicts() ──
#> ✖ dplyr::filter() masks stats::filter()
#> ✖ dplyr::lag()    masks stats::lag()
#> ℹ Use the conflicted package (<http://conflicted.r-lib.org/>) to force all conflicts to become errors
library("tximport")
library("biomaRt")
library("DESeq2")
#> Loading required package: S4Vectors
#> Loading required package: stats4
#> Loading required package: BiocGenerics
#>
#> Attaching package: 'BiocGenerics'
#>
#> The following objects are masked from 'package:lubridate':
#>
#>     intersect, setdiff, union
#>
#> The following objects are masked from 'package:dplyr':
#>
#>     combine, intersect, setdiff, union
#>
#> The following objects are masked from 'package:stats':
#>
#>     IQR, mad, sd, var, xtabs
#>
#> The following objects are masked from 'package:base':
#>
#>     anyDuplicated, aperm, append, as.data.frame, basename, cbind,
#>     colnames, dirname, do.call, duplicated, eval, evalq, Filter, Find,
#>     get, grep, grepl, intersect, is.unsorted, lapply, Map, mapply,
#>     match, mget, order, paste, pmax, pmax.int, pmin, pmin.int,
#>     Position, rank, rbind, Reduce, rownames, sapply, setdiff, sort,
#>     table, tapply, union, unique, unsplit, which.max, which.min
#>
#>
#> Attaching package: 'S4Vectors'
#>
#> The following objects are masked from 'package:lubridate':
#>
#>     second, second<-
#>
#> The following objects are masked from 'package:dplyr':
#>
#>     first, rename
#>
#> The following object is masked from 'package:tidyr':
#>
#>     expand
#>
#> The following object is masked from 'package:utils':
#>
#>     findMatches
#>
#> The following objects are masked from 'package:base':
#>
#>     expand.grid, I, unname
#>
#> Loading required package: IRanges
#>
#> Attaching package: 'IRanges'
#>
#> The following object is masked from 'package:lubridate':
#>
#>     %within%
#>
#> The following objects are masked from 'package:dplyr':
#>
#>     collapse, desc, slice
#>
#> The following object is masked from 'package:purrr':
#>
#>     reduce
#>
#> Loading required package: GenomicRanges
#> Loading required package: GenomeInfoDb
#> Loading required package: SummarizedExperiment
#> Loading required package: MatrixGenerics
#> Loading required package: matrixStats
#>
#> Attaching package: 'matrixStats'
#>
#> The following object is masked from 'package:dplyr':
#>
#>     count
#>
#>
#> Attaching package: 'MatrixGenerics'
#>
#> The following objects are masked from 'package:matrixStats':
#>
#>     colAlls, colAnyNAs, colAnys, colAvgsPerRowSet, colCollapse,
#>     colCounts, colCummaxs, colCummins, colCumprods, colCumsums,
#>     colDiffs, colIQRDiffs, colIQRs, colLogSumExps, colMadDiffs,
#>     colMads, colMaxs, colMeans2, colMedians, colMins, colOrderStats,
#>     colProds, colQuantiles, colRanges, colRanks, colSdDiffs, colSds,
#>     colSums2, colTabulates, colVarDiffs, colVars, colWeightedMads,
#>     colWeightedMeans, colWeightedMedians, colWeightedSds,
#>     colWeightedVars, rowAlls, rowAnyNAs, rowAnys, rowAvgsPerColSet,
#>     rowCollapse, rowCounts, rowCummaxs, rowCummins, rowCumprods,
#>     rowCumsums, rowDiffs, rowIQRDiffs, rowIQRs, rowLogSumExps,
#>     rowMadDiffs, rowMads, rowMaxs, rowMeans2, rowMedians, rowMins,
#>     rowOrderStats, rowProds, rowQuantiles, rowRanges, rowRanks,
#>     rowSdDiffs, rowSds, rowSums2, rowTabulates, rowVarDiffs, rowVars,
#>     rowWeightedMads, rowWeightedMeans, rowWeightedMedians,
#>     rowWeightedSds, rowWeightedVars
#>
#> Loading required package: Biobase
#> Welcome to Bioconductor
#>
#>     Vignettes contain introductory material; view with
#>     'browseVignettes()'. To cite Bioconductor, see
#>     'citation("Biobase")', and for packages 'citation("pkgname")'.
#>
#>
#> Attaching package: 'Biobase'
#>
#> The following object is masked from 'package:MatrixGenerics':
#>
#>     rowMedians
#>
#> The following objects are masked from 'package:matrixStats':
#>
#>     anyMissing, rowMedians
library("GenomicRanges")
library("PCAtools")
#> Loading required package: ggrepel
#>
#> Attaching package: 'PCAtools'
#>
#> The following objects are masked from 'package:stats':
#>
#>     biplot, screeplot
library("ComplexHeatmap")
#> Loading required package: grid
#> ========================================
#> ComplexHeatmap version 2.18.0
#> Bioconductor page: http://bioconductor.org/packages/ComplexHeatmap/
#> Github page: https://github.com/jokergoo/ComplexHeatmap
#> Documentation: http://jokergoo.github.io/ComplexHeatmap-reference
#>
#> If you use it in published research, please cite either one:
#> - Gu, Z. Complex Heatmap Visualization. iMeta 2022.
#> - Gu, Z. Complex heatmaps reveal patterns and correlations in multidimensional
#>     genomic data. Bioinformatics 2016.
#>
#>
#> The new InteractiveComplexHeatmap package can directly export static
#> complex heatmaps into an interactive Shiny app with zero effort. Have a try!
#>
#> This message can be suppressed by:
#>   suppressPackageStartupMessages(library(ComplexHeatmap))
#> ========================================
library("fgsea")
library("msigdbr")
library("eulerr")
library("patchwork")
```

#### Prepare Data

Salmon estimates *transcript* level abundance. To summarize to *gene* level abundance a transcript to gene gene mapping `data.frame` must be created. Biomart will be queried for this.

Hide

```
ensembl <- useEnsembl(
  biomart="genes", dataset="dmelanogaster_gene_ensembl", version=110
)

tx2gene <- getBM(
  attributes=c("ensembl_transcript_id", "ensembl_gene_id"),
  mart=ensembl
)
```

After the transcript to gene map is created, `tximport` can be used to load the transcript abundance estimates and summarize to the gene level.

Hide

```
quant_files <-
  file.path("resources", "quants") |>
  list.files(recursive=TRUE, pattern="quant.sf", full.names=TRUE)

names(quant_files) <- str_extract(
  quant_files,
  "(?<=/quants/)[^/]+(?=/quant\\.sf)"
)

quant_mat <- tximport(
  quant_files, type="salmon", tx2gene=tx2gene, dropInfReps=TRUE
)
#> reading in files with read_tsv
#> 1 2 3 4 5 6 7 8 9 10 11 12
#> summarizing abundance
#> summarizing counts
#> summarizing length
```

Load the metadata `csv` file which contains supplemental information about the samples that is useful for downstream analysis.

Hide

```
metadata <-
  file.path("resources", "sample_sheet.csv") |>
  read_csv() |>
  dplyr::mutate(
    source=factor(source, levels=c("whole_body", "fat_body")),
    condition=factor(condition, levels=c("control", "err_mut"))
  ) |>
  column_to_rownames("sample_name")
#> Rows: 12 Columns: 3
#> ── Column specification ────────────────────────────────────────────
#> Delimiter: ","
#> chr (3): sample_name, source, condition
#>
#> ℹ Use `spec()` to retrieve the full column specification for this data.
#> ℹ Specify the column types or set `show_col_types = FALSE` to quiet this message.

# Ensure the samples are in the same order as the quantification matrix.
metadata <- metadata[colnames(quant_mat$counts), ]
```

Create a DESeq2 object using the gene-level abundance estimates along with the sample metadata.

The two effects that are of interest to us are the **condition** which represents the mutant status of ERR, and **source** which is either the ‘whole body’ or ‘fat body’.

This model also contains an interaction term `condition:source` that captures the effect sample source has on the response to ERR mutant status. Another way of thinking about is is that it checks whether the difference between the ERR mutant and control is different in whole body versus fat body.

The contrast can be thought of as the following:

$$|(W_m - W_c) - (F_m - F_c)| > 0$$

Where $W_m$ and $W_c$ are the whole body mutant and control, and $F_m$ and $F_c$ are the fat body mutant and control.

Hide

```
dds <- DESeqDataSetFromTximport(
  quant_mat, metadata, ~ condition + source + condition:source
)
#> using counts and average transcript lengths from tximport
```

Once the DESeq2 object is created, Biomart is queried to add more useful information for each gene such as their genomic coordinates and biotype (protein coding, lncRNA, etc.).

Hide

```
gene_info <-
  getBM(
    attributes=c(
      "ensembl_gene_id", "external_gene_name",
      "description", "source", "gene_biotype",
      "chromosome_name", "start_position", "end_position", "strand"
    ),
    mart=ensembl
  ) |>
  dplyr::rename(
    seqnames="chromosome_name", start="start_position", end="end_position"
  ) |>
  mutate(
    strand=ifelse(strand == -1, "-", "+"),
    across(where(is.character), \(x) na_if(x, ""))
  )

rownames(gene_info) <- gene_info$ensembl_gene_id
gene_info <- makeGRangesFromDataFrame(gene_info, keep.extra.columns=TRUE)

rowRanges(dds) <- gene_info[rownames(dds), ]
```

#### Count Normalization

For various downstream purposes such as PCA the counts must be normalized. `rlog` transformation normalizes counts using variance stabalization (to reduce the mean expression-variance correlation) and preserving the effect differences specified in our model (via `blind=FALSE`). The normalized counts are saved back into the DESeq2 object.

Hide

```
assay(dds, "rlog") <- assay(rlog(dds, blind=FALSE))
#> using 'avgTxLength' from assays(dds), correcting for library size
```

#### PCA Analysis

To diagnose any postential sample problems and as a preliminary exploration of variance sources in the dataset PCA anlysis is performed. The rlog counts, centered, are used as input.

Hide

```
pca_results <- pca(
  assay(dds, "rlog"), removeVar=0.1, metadata=colData(dds),
  center=TRUE, scale=FALSE
)
#> -- removing the lower 10% of variables based on variance
```

A scree plot allows one to explore how much variance each principle component captures, along with the cumulative sum of explained variance over progessive components.

In the below scree plot `PC1` captures a majority of the variance (~85%).

Hide

```
p <- screeplot(pca_results)
print(p)
```

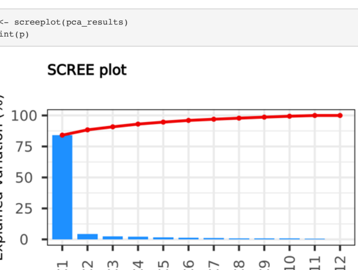


The scree plot shows that ~90% of variance is explained by the first two principle coponents. A biplot of these components is created with point color as ERR mutant status, and point shape as tissue source.

It is clear when looking at the below biplot that the majority of the variance (~85%) in the dataset is associated with the sample tissue source, which is expected. The second most variance (~4%) captures the difference between the ERR mutants.

Overall the PCA plot shows promising results, with the first to principle components capturing the main effects being studied in the experiment, and the samples grouping well into each condition.

Hide

```
p <- biplot(pca_results, colby="condition", shape="source")
print(p)
```

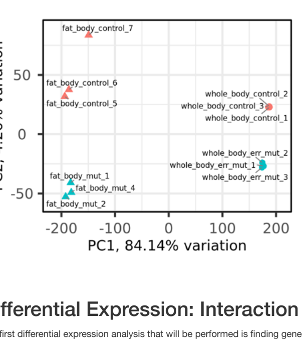


#### Differential Expression: Interaction Term.

The first differential expression analysis that will be performed is finding genes with a significant interaction term.

As previously discussed, an interaction term is used to capture genes where the difference between the ERR mutant and control is different in whole body versus fat body.

Hide

```
dds <- DESeq(dds)
#> estimating size factors
#> using 'avgTxLength' from assays(dds), correcting for library size
#> estimating dispersions
#> gene-wise dispersion estimates
#> mean-dispersion relationship
#> final dispersion estimates
#> fitting model and testing
```

After the differential expression above, the results are returned and filtered for genes with significant interaction terms.

Hide

```
results_interaction <- dds |>
  results(name="conditionerr_mut.sourcefat_body") |>
  as_tibble(rownames="ensembl_gene_id") |>
  left_join(
    dds |>
      rowData() |>
      as_tibble() |>
      dplyr::select(ensembl_gene_id, external_gene_name, gene_biotype),
    by="ensembl_gene_id"
  ) |>
  dplyr::select(
    ensembl_gene_id, external_gene_name, gene_biotype,
    baseMean, stat, pvalue, padj
  ) |>
  arrange(padj)
```

| ensembl_gene_id | external_gene_name | gene_biotype | baseMean | stat | pvalue | padj |
|---|---|---|---|---|---|---|
| FBgn0036824 | CG3902 | protein_coding | 8006.0021 | -22.47023 | 0 | 0 |
| FBgn0263782 | Hmgcr | protein_coding | 1221.8905 | 20.44253 | 0 | 0 |
| FBgn0016123 | Alp4 | protein_coding | 2864.8304 | 17.71378 | 0 | 0 |
| FBgn0023537 | CG17896 | protein_coding | 12103.6785 | -17.16675 | 0 | 0 |
| FBgn0261555 | CG42673 | protein_coding | 487.0214 | 15.94050 | 0 | 0 |
| FBgn0034406 | Jheh3 | protein_coding | 1662.4675 | -15.62885 | 0 | 0 |
| FBgn0030612 | Dbct | protein_coding | 1174.9950 | -14.63281 | 0 | 0 |
| FBgn0039678 | Obp99a | protein_coding | 5222.9189 | -13.70555 | 0 | 0 |
| FBgn0010482 | hlk | protein_coding | 4606.2367 | 13.65649 | 0 | 0 |
| FBgn0040064 | yip2 | protein_coding | 3956.2902 | -13.51764 | 0 | 0 |

The above table shows the top 10 of the roughly 1.3k genes with a significant interaction terms for illustrative purposes. By themselves these results are somewhat difficult to interpret because although it returns significant interactions, no information is returned for the direction and magnitude of this difference.

To rectify this, the analyses that follow will look for genes with a statistically significant difference between ERR and control in the whole body, and separatly in the fat body. After these two contrast only genes with a statistically significant interaction term will be retained.

#### Differential Expression: Mutants vs. Controls.

As described in the interaction term section, differential expression will be be performed to compare both the whole body ERR mutant vs. control, and separatly the fat body ERR mutant vs. control. These contrasts will be explored first without consideration of the interaction term, and then later differentially expressed genes for these comparisons will be filtered by whether the gene had a significant interaction term.

The first comparison will be the whole body ERR mutant vs. control. The returned results will both have their $log_2$ FC shrunk (to compensate for the mean expression-$log_2$FC corelation) and will be tested against $|log_2 FC| > 1$ using the `apeglm` library.

Testing directly against a $log_2$ FC threshold is advantageous versus a nieve filtering approach because it introduces statistical rigor via considering the uncertainty of the computed $log_2$ FC when determining whether the gene is beyond the target threshold. The `apeglm` library will return an $s$-value for this threshold test, which represents the confidence that a change is of a certain size. Typically a lower threshold than $FDR$ is used $s$-value $< 0.005$ because an $s$-value of 0.5 corresponds to random guessing whereas an $FDR$ of 1 corresponds to all nulls.

Hide

```
# Create a 'group' factor that concatenates the condition with the source.
# This allows whole body ERR mutant vs. whole body control.
dds[["group"]] <- factor(
  str_glue("{dds[['source']]}_{dds[['condition']]}"),
  levels=c(
    "whole_body_control", "whole_body_err_mut",
    "fat_body_control", "fat_body_err_mut"
  )
)

design(dds) <- ~ group
dds <- DESeq(dds)
#> using pre-existing normalization factors
#> estimating dispersions
#> found already estimated dispersions, replacing these
#> gene-wise dispersion estimates
#> mean-dispersion relationship
#> final dispersion estimates
#> fitting model and testing

# Return the results with log2FC shrinkage and testing whether the
# absolute value of the log2FC is greater than 1.
results_wb <- dds |>
  lfcShrink(
    coef="group_whole_body_err_mut_vs_whole_body_control", lfcThreshold=1
  ) |>
  as_tibble(rownames="ensembl_gene_id") |>
  left_join(
    rowData(dds) |>
      as_tibble() |>
      dplyr::select(ensembl_gene_id, external_gene_name, gene_biotype),
    by="ensembl_gene_id"
  ) |>
  dplyr::mutate(contrast="whole_body_err_mut_vs_control") |>
  dplyr::relocate(external_gene_name, gene_biotype, .after=ensembl_gene_id)
#> using 'apeglm' for LFC shrinkage. If used in published research, please cite:
#>     Zhu, A., Ibrahim, J.G., Love, M.I. (2018) Heavy-tailed prior distributions for
#>     sequence count data: removing the noise and preserving large differences.
#>     Bioinformatics. https://doi.org/10.1093/bioinformatics/bty895
#> computing FSOS 'false sign or small' s-values (T=1)
```

A table showing the top 10 differentially expressed genes by $s$-value and $|log_2 \text{FC}|$ for the whole body ERR mutant vs. control comparison.

| ensembl_gene_id | external_gene_name | gene_biotype | baseMean | log2FoldChange | lfcSE | svalue |
|---|---|---|---|---|---|---|
| FBgn0034710 | Alp7 | protein_coding | 127.71808 | -9.726892 | 1.0056687 | 0 |
| FBgn0030459 | CG12723 | protein_coding | 92.44591 | -8.006515 | 0.8136604 | 0 |
| FBgn0034712 | Alp8 | protein_coding | 160.96855 | -6.847773 | 0.5116262 | 0 |
| FBgn0029681 | CG15239 | protein_coding | 263.78509 | -6.130804 | 0.3847080 | 0 |
| FBgn0001258 | Ldh | protein_coding | 1260.22435 | -5.518817 | 0.1570752 | 0 |
| FBgn0043783 | CG32444 | protein_coding | 421.79447 | -5.502725 | 0.3667243 | 0 |
| FBgn0036350 | CG14111 | protein_coding | 107.81346 | -4.817017 | 0.3528751 | 0 |
| FBgn0034391 | CG15080 | protein_coding | 181.67706 | -4.810925 | 0.4014001 | 0 |
| FBgn0263038 | CG43333 | protein_coding | 204.31932 | -4.689629 | 0.2991950 | 0 |
| FBgn0259233 | CG42331 | protein_coding | 99.00688 | -4.540791 | 0.4158427 | 0 |

Volcano plots are a useful visualization of the results of differential expression. The $log_2$ FC is plotted against the $-log_{10}(s\text{-value})$ where the $s$-value is returned from `apeglm` after $log_2$ Fold Change shrinkage and testing against $|log_2 \text{ Fold Change}| > 1$.

Points with an $s$-value $< 0.005$ are colored red, and dotted lines are drawn at $s$-value $= 0.005$ and $log_2$ Fold Change $= \pm 1$. A dashed line is drawn at $log_2$ Fold Change $= 1$ as an easy demarcation of genes which are up and down regulated.

Hide

```
results_wb |>
  dplyr::mutate(
    deg_status=ifelse(
      svalue > 0.005 | is.na(svalue),
      "svalue > 0.005 (|log2FC| > 1)",
      "svalue < 0.005 (|log2FC| < 1)"
    )
  ) |>
  ggplot(aes(x=log2FoldChange, y=-log10(svalue))) +
    geom_point(aes(color=deg_status), size=0.5) +
    theme_bw() +
    scale_color_manual(values=c("salmon", "lightgrey")) +
    geom_vline(xintercept=0, linetype="dashed", size=0.5, color="dimgrey") +
    geom_vline(xintercept=1, linetype="dotted", size=0.5, color="dimgrey") +
    geom_vline(xintercept=-1, linetype="dotted", size=0.5, color="dimgrey") +
    geom_hline(
      yintercept=-log10(0.005), linetype="dotted", size=0.5, color="dimgrey"
    ) +
    guides(color=guide_legend(override.aes=list(size=2), ncol=1)) +
    labs(
      x=bquote(log[2]~Fold~Change),
      y=bquote(-log[10](italic(s)*-value)),
      color="Differential\nStatus",
      title="Whole Body ERR Mutant\nDifferentially Expressed Genes"
    )+
    theme(
      legend.position="bottom",
      title=element_text(size=8)
    )
#> Warning: Using `size` aesthetic for lines was deprecated in ggplot2 3.4.0.
#> ℹ Please use `linewidth` instead.
#> This warning is displayed once every 8 hours.
#> Call `lifecycle::last_lifecycle_warnings()` to see where this warning was generated.
#> Warning: Removed 10001 rows containing missing values (`geom_point()`).
```

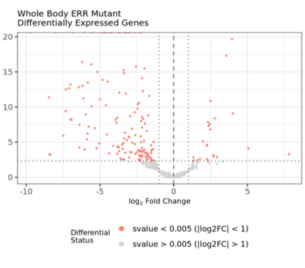


The second comparison is the fat body ERR mutant vs. control and the same analysis steps as described for the whole body comparison will be performed.

Hide

```
# Create a 'group' factor that concatenates the condition with the source.
# This allows fat body ERR mutant vs. fat body control.
dds[["group"]] <- factor(
  dds[["group"]],
  levels=c(
    "fat_body_control", "fat_body_err_mut",
    "whole_body_control", "whole_body_err_mut"
  )
)

design(dds) <- ~ group
dds <- DESeq(dds)
#> using pre-existing normalization factors
#> estimating dispersions
#> found already estimated dispersions, replacing these
#> gene-wise dispersion estimates
#> mean-dispersion relationship
#> final dispersion estimates
#> fitting model and testing

# Return the results with log2FC shrinkage and testing whether the
# absolute value of the log2FC is greater than 1.
results_fb <- dds |>
  lfcShrink(
    coef="group_fat_body_err_mut_vs_fat_body_control", lfcThreshold=1
  ) |>
  as_tibble(rownames="ensembl_gene_id") |>
  left_join(
    rowData(dds) |>
      as_tibble() |>
      dplyr::select(ensembl_gene_id, external_gene_name, gene_biotype),
    by="ensembl_gene_id"
  ) |>
  dplyr::mutate(contrast="fat_body_err_mut_vs_control") |>
  dplyr::relocate(external_gene_name, gene_biotype, .after=ensembl_gene_id)
#> using 'apeglm' for LFC shrinkage. If used in published research, please cite:
#>     Zhu, A., Ibrahim, J.G., Love, M.I. (2018) Heavy-tailed prior distributions for
#>     sequence count data: removing the noise and preserving large differences.
#>     Bioinformatics. https://doi.org/10.1093/bioinformatics/bty895
#> computing FSOS 'false sign or small' s-values (T=1)
```

A table showing the top 10 differentially expressed genes by $s$-value and $|log_2 \text{FC}|$ for the fat body ERR mutant vs. control comparison.

| ensembl_gene_id | external_gene_name | gene_biotype | baseMean | log2FoldChange | lfcSE | svalue |
|---|---|---|---|---|---|---|
| FBgn0024289 | Sodh-1 | protein_coding | 3728.6176 | -7.445477 | 0.2856145 | 0 |
| FBgn0032945 | CG8665 | protein_coding | 1242.8646 | -7.198340 | 0.4107776 | 0 |
| FBgn0043783 | CG32444 | protein_coding | 421.7945 | -6.882986 | 0.4306172 | 0 |
| FBgn0039678 | Obp99a | protein_coding | 5222.9189 | -6.808595 | 0.2725226 | 0 |
| FBgn0027601 | pdgy | protein_coding | 767.5290 | -6.785336 | 0.4858414 | 0 |
| FBgn0085195 | CG34166 | protein_coding | 8371.0366 | -6.369281 | 0.3631577 | 0 |
| FBgn0034356 | Pepck2 | protein_coding | 1107.9962 | -6.160579 | 0.3148432 | 0 |
| FBgn0286508 | Had1 | protein_coding | 3161.4824 | -5.110121 | 0.2828146 | 0 |
| FBgn0086355 | Tpi | protein_coding | 2840.3978 | -4.869012 | 0.1583299 | 0 |
| FBgn0003074 | Pgi | protein_coding | 5791.1554 | -4.560174 | 0.1431879 | 0 |

Once again a volcano plot is generated to visualize the differentially expressed genes for the fat body ERR mutant vs. control comparison.

Hide

```
results_fb |>
  dplyr::mutate(
    deg_status=ifelse(
      svalue > 0.005 | is.na(svalue),
      "svalue > 0.005 (|log2FC| > 1)",
      "svalue < 0.005 (|log2FC| < 1)"
    )
  ) |>
  ggplot(aes(x=log2FoldChange, y=-log10(svalue))) +
    geom_point(aes(color=deg_status), size=0.5) +
    theme_bw() +
    scale_color_manual(values=c("salmon", "lightgrey")) +
    geom_vline(xintercept=0, linetype="dashed", size=0.5, color="dimgrey") +
    geom_vline(xintercept=1, linetype="dotted", size=0.5, color="dimgrey") +
    geom_vline(xintercept=-1, linetype="dotted", size=0.5, color="dimgrey") +
    geom_hline(
      yintercept=-log10(0.005), linetype="dotted", size=0.5, color="dimgrey"
    ) +
    guides(color=guide_legend(override.aes=list(size=2), ncol=1)) +
    labs(
      x=bquote(log[2]~Fold~Change),
      y=bquote(-log[10](italic(s)*-value)),
      color="Differential\nStatus",
      title="Fat Body ERR Mutant\nDifferentially Expressed Genes"
    )+
    theme(
      legend.position="bottom",
      title=element_text(size=8)
    )
#> Warning: Removed 10001 rows containing missing values (`geom_point()`).
```

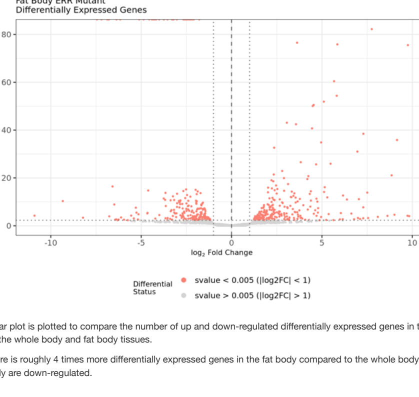


A bar plot is plotted to compare the number of up and down-regulated differentially expressed genes in the ERR mutant vs. control comparison for the whole body and fat body tissues.

There is roughly 4 times more differentially expressed genes in the fat body compared to the whole body, and most differential genes in the whole body are down-regulated.

Hide

```
p <- bind_rows(results_wb, results_fb) |>
  dplyr::filter(svalue < 0.005) |>
  dplyr::mutate(
    deg_status=ifelse(log2FoldChange > 0, "Up", "Down"),
    deg_status=factor(deg_status, levels=c("Up", "Down")),
    group=case_when(
      contrast == "fat_body_err_mut_vs_control" ~ "Fat Body",
      contrast == "whole_body_err_mut_vs_control" ~ "Whole Body"
    ),
    group=factor(group, levels=c("Whole Body", "Fat Body"))
  ) |>
  dplyr::count(contrast, group, deg_status) |>
  ggplot(aes(x=group, y=n, fill=deg_status)) +
    geom_col(width=0.5) +
    theme_bw() +
    theme(
      axis.text.x=element_text(angle=45, hjust=1),
      legend.position="top",
      plot.title.position="plot",
      axis.title.x=element_blank()
    ) +
    labs(
      y="Gene Count", x="Comparison", fill="Differential\nStatus:",
      title="Differentially Expressed Genes:\nWhole Body vs Fat Body"
    ) +
    scale_fill_manual(values=c("Down"="salmon", "Up"="darkturquoise"))

print(p)
```

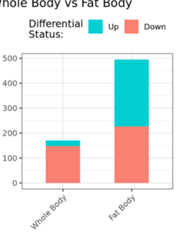


Hide

```
ggsave(
  file.path("results", "degs_barplot_whole_body_vs_fat_body.pdf"), plot=p,
  device=cairo_pdf, height=4, width=3
)
```

A euler plot is a venn diagram but with the circle sizes and degree of overlap scaled proportionally to the number of values in each group. I will generate a euler plot that considers separatly the up and down-regulated genes in ERR mutants vs. control groups for the whole body and fat body.

```r
euler_list <- bind_rows(results_wb, results_fb) |>
  dplyr::filter(svalue < 0.005) |>
  dplyr::mutate(
    deg_status=ifelse(log2FoldChange > 0, "Up", "Down"),
    group=case_when(
      contrast == "fat_body_err_mut_vs_control" ~ "Fat Body",
      contrast == "whole_body_err_mut_vs_control" ~ "Whole Body"
    ),
    label=str_glue("{group} {deg_status}")
  ) |>
  split(~ label) |>
  map(pull, ensembl_gene_id)

p <- euler(euler_list, shape="circle") |>
  plot(
    fill=c("deeppink", "orange", "turquoise", "yellowgreen"), col="black",
    lwd=1.5, alpha=1, quantities=list(cex=1), labels=NULL,
    legend=list(labels=names(euler_list)),
    main=list(
      label="Overlap of Differential Genes:\nWhole Body vs. Fat Body",
      cex=1.15
    )
  )

print(p)
```

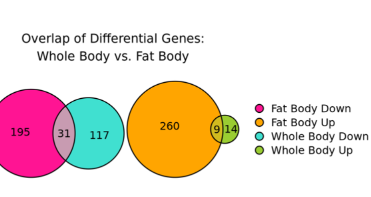


Term enrichment will be the final analysis of the separate whole body and fat body comparisons. The genes that are up or down-regulated in either of the tissues will be tested against terms aggregated by the Molecular Signatures Database (MSigDB) using a hypergeometric test of enrichment. More specifically, the following collections will be tested:

- **Hallmark Gene Sets**: Curated MSigDB signatures.
- **Reactome Pathways**: Pathways from the reactome database.
- **Chemical and Genetic Perturbations**: Gene sets from papers testing chemical or genetic perturbations.
- **Gene Ontology Molecular Function**: The molecular-level activities of the gene product.
- **Gene Ontology Biological Process**: The larger biological processes the gene product is involved in.
- **Human Phenotype Ontology**: Gene sets characterized into groups corresponding to a standardized vocabulary of phenotypic human abnormalities.

Refer to the linked website above for more information on each of the collections.

```r
deg_list <- bind_rows(results_wb, results_fb) |>
  dplyr::filter(svalue < 0.005) |>
  dplyr::mutate(
    deg_status=ifelse(log2FoldChange > 0, "Up", "Down"),
    group=case_when(
      contrast == "fat_body_err_mut_vs_control" ~ "Fat Body",
      contrast == "whole_body_err_mut_vs_control" ~ "Whole Body"
    ),
    label=str_glue("{group} {deg_status}")
  ) |>
  split(~ label) |>
  map(pull, ensembl_gene_id)

term_enrichment <- function(degs, all_genes, category="H", subcategory="") {
  geneset <-
    msigdbr(
      species="Drosophila melanogaster",
      category=category, subcategory=subcategory
    ) |>
    dplyr::select(ensembl_gene, gs_name) |>
    split(~ gs_name) |>
    map(pull, ensembl_gene)

  universe <- union(
    unique(unname(unlist(geneset))),
    all_genes
  )

  enrich <-
    fora(pathways=geneset, genes=degs, universe=universe) |>
    as_tibble()

  return(enrich)
}

enrich_results <- list()

enrich_results[["Hallmark Gene Sets"]] <- map(
  deg_list, term_enrichment,
  rownames(dds), category="H", subcategory=""
)

enrich_results[["Reactome Pathways"]] <- map(
  deg_list, term_enrichment,
  rownames(dds), category="C2", subcategory="CP:REACTOME"
)

enrich_results[["Chemical and Genetic Perturbations"]] <- map(
  deg_list, term_enrichment,
  rownames(dds), category="C2", subcategory="CGP"
)

enrich_results[["Gene Ontology Molecular Function"]] <- map(
  deg_list, term_enrichment,
  rownames(dds), category="C5", subcategory="GO:MF"
)

enrich_results[["Gene Ontology Biological Process"]] <- map(
  deg_list, term_enrichment,
  rownames(dds), category="C5", subcategory="GO:BP"
)

enrich_results[["Human Phenotype Ontology"]] <- map(
  deg_list, term_enrichment,
  rownames(dds), category="C5", subcategory="HPO"
)
```

A dot plot will be used to help visualize the enrichment results. Gene sets are filted via $FDR < 0.05$ and for each comparison up to the top 10 enriched terms (by lowest $FDR$) are retained. The x-axis is the $-log_{10}(FDR)$, the point colors are whether $FDR < 0.05$ and the point sizes correspond to the fraction of genes in the differential expression list for that term versus the total number of genes in that term. The terms are ordered on the y-axis according to the minimum $FDR$ value osberved among all comparisons.

```r
p <- plot_enrichment <- function(enrich_results, y_lab="Term") {
  # Get at most the top 10 pathways by lowest FDR for each list element.
  top_pathways <- enrich_results |>
    bind_rows(.id="group") |>
    dplyr::filter(padj < 0.05) |>
    dplyr::group_by(group) |>
    dplyr::slice_min(padj, n=10) |>
    pull(pathway) |>
    unique()

  # Generate the plot.
  p <- enrich_results |>
    bind_rows(.id="group") |>
    dplyr::filter(pathway %in% top_pathways) |>
    dplyr::mutate(
    frac= overlap / size,
    pathway=pathway |>
      str_remove("^(REACTOME|HP|HALLMARK|GOMF|GOP)_") |>
      str_replace_all("_", " "),
    pathway=fct_reorder(pathway, -log10(padj), .fun=median),
    sig=ifelse(padj < 0.05, "FDR < 0.05", "FDR > 0.05")
    ) |>
    ggplot(aes(x=-log10(padj), y=pathway)) +
    geom_segment(aes(xend=0, yend=pathway), linewidth=0.25, color="dimgrey") +
    geom_point(aes(size=frac, color=sig)) +
    facet_grid(cols=vars(group)) +
    theme_bw() +
    labs(
      x=bquote(-log[10](FDR)), y=y_lab,
      color="Enrichment\nSignificance",
      size="Fraction of Genes\nin Pathway"
    ) +
    theme(
      legend.position="right", legend.box="vertical",
      axis.text.y=element_text(size=7.5),
      axis.text.x=element_text(angle=45, hjust=1)
    ) +
    guides(color=guide_legend(override.aes=list(size=2.5)))

  return(p)
}

p <- imap(enrich_results, plot_enrichment)

p <- wrap_plots(p, ncol=1)

print(p)
```

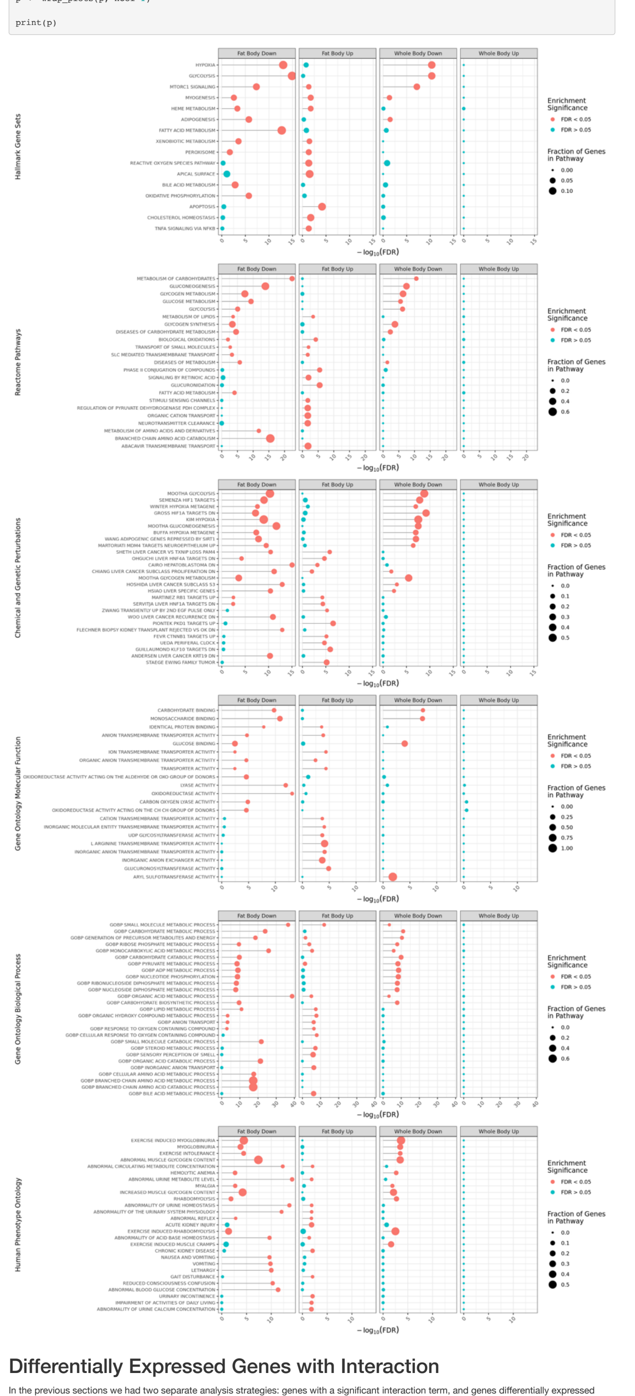

#### Differentially Expressed Genes with Interaction

In the previous sections we had two separate analysis strategies: genes with a significant interaction term, and genes differentially expressed between the ERR mutant and control in either the whole body or fat body. In this section we will combine these analysis by finding genes that are both differentially expressed *and* have a significant interaction term.

Intuitively this means we are looking for genes that are differentially expressed in their respective tissues, and whose $log2FC$ is statistically different in the fat body versus the whole body (or vice versa). Here are some illustrative examples of how this could look in practice in the data:

- A gene that is not differentially expressed in the whole body, but down-regulated in the fat body.
- A gene that is slightly down-regulated in the fat body, but greatly down-regulated in the whole body.
- A gene that is up-regulated in the whole body, but down-regulated in the fat body.
- …

To get an overall idea of how many of the differentially expressed genes have an interaction term, a euler plot is generated.

```r
euler_list <- bind_rows(results_wb, results_fb) |>
  dplyr::filter(svalue < 0.005) |>
  dplyr::mutate(
    deg_status=ifelse(log2FoldChange > 0, "Up", "Down"),
    group=case_when(
      contrast == "fat_body_err_mut_vs_control" ~ "Fat Body",
      contrast == "whole_body_err_mut_vs_control" ~ "Whole Body"
    ),
    label=str_glue("{group} {deg_status}")
  ) |>
  split(~ label) |>
  map(pull, ensembl_gene_id)

euler_list[["Significant Interaction"]] <- results_interaction |>
  dplyr::filter(padj < 0.05) |>
  pull(ensembl_gene_id)

p <- euler(euler_list, shape="ellipse") |>
  plot(
    fill=c("deeppink", "orange", "turquoise", "yellowgreen", "lightgrey"),
    col="black", lwd=1.5, alpha=1, quantities=list(cex=1), labels=NULL,
    legend=list(labels=names(euler_list)),
    main=list(
      label="Differential Genes With Interactions",
      cex=1.15
    )
  )

print(p)
```

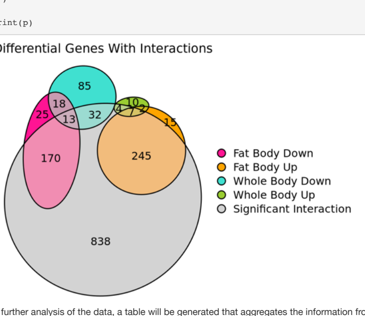


For further analysis of the data, a table will be generated that aggregates the information from the differential expression analysis and the interaction terms for all genes.

Genes with an $s$-value $< 0.005$ and a $|log2\,\mathrm{FC}| > 1$ are marked as 'Up' or 'Down', and genes with $s$-value $> 0.005$ are marked as 'n.s.'.

```r
degs_and_interactions <-
  full_join(
    results_fb |>
      dplyr::select(
        ensembl_gene_id, external_gene_name,
        fb_log2FoldChange=log2FoldChange, fb_svalue=svalue,
        gene_biotype
      ) |>
      dplyr::mutate(
        fb_status=case_when(
          fb_svalue < 0.005 & fb_log2FoldChange > 0 ~ "Up",
          fb_svalue < 0.005 & fb_log2FoldChange < 0 ~ "Down",
          TRUE ~ "n.s."
        )
      ),
    results_wb |>
      dplyr::select(
        ensembl_gene_id, external_gene_name,
        wb_log2FoldChange=log2FoldChange, wb_svalue=svalue,
        gene_biotype
      ) |>
      dplyr::mutate(
        wb_status=case_when(
          wb_svalue < 0.005 & wb_log2FoldChange > 0 ~ "Up",
          wb_svalue < 0.005 & wb_log2FoldChange < 0 ~ "Down",
          TRUE ~ "n.s."
        )
      )
  ) |>
  left_join(
    dplyr::select(results_interaction, ensembl_gene_id, interaction_padj=padj),
    by="ensembl_gene_id"
  )
#> Joining with `by = join_by(ensembl_gene_id, external_gene_name, gene_biotype)`
```

A table of the top 10 differentially expressed genes by interaction term $FDR$.

| ensembl_gene_id | external_gene_name | fb_log2FoldChange | fb_svalue | gene_biotype | fb_status | wb_log2FoldChange | wb_svalue | wb_status | interaction_padj |
|---|---|---|---|---|---|---|---|---|---|
| FBgn0036824 | CG3902 | -2.570618 | 0 | protein_coding | Down | -0.1166981 | 0.9771123 | n.s. | 0 |
| FBgn0263782 | Hmgcr | 3.623422 | 0 | protein_coding | Up | -0.3713812 | 0.8350958 | n.s. | 0 |
| FBgn0016123 | Alp4 | 9.754924 | 0 | protein_coding | Up | 0.0057770 | 0.9677967 | n.s. | 0 |
| FBgn0023537 | CG17896 | -2.082266 | 0 | protein_coding | Down | -0.0365163 | 0.9760630 | n.s. | 0 |
| FBgn0261555 | CG42673 | 5.851735 | 0 | protein_coding | Up | 0.5056880 | 0.4985521 | n.s. | 0 |
| FBgn0034406 | Jheh3 | -4.256450 | 0 | protein_coding | Down | 0.0468420 | 0.9711870 | n.s. | 0 |
| FBgn0030612 | Dbct | -1.790013 | 0 | protein_coding | Down | -0.0483963 | 0.9764596 | n.s. | 0 |
| FBgn0039678 | Obp99a | -6.808595 | 0 | protein_coding | Down | -1.5597686 | 0.0018714 | Down | 0 |
| FBgn0010482 | hlk | 5.105570 | 0 | protein_coding | Up | -0.0053842 | 0.9683309 | n.s. | 0 |
| FBgn0040064 | yip2 | -4.347433 | 0 | protein_coding | Down | -0.6134745 | 0.4316501 | n.s. | 0 |

A bar plot of differentially expressed genes is once again generated, but this time only genes with a significant interaction term are retained and counted.

```r
p <- degs_and_interactions |>
  dplyr::filter(
    (fb_svalue < 0.005 | wb_svalue < 0.005) & interaction_padj < 0.05
  ) |>
  dplyr::count(fb_status, wb_status) |>
  dplyr::filter(!(fb_status == "n.s." & wb_status == "n.s.")) |>
  pivot_longer(!n, names_to="tissue", values_to="deg_status") |>
  dplyr::group_by(tissue, deg_status) |>
  dplyr::summarize(n=sum(n), .groups="drop") |>
  dplyr::filter(deg_status != "n.s.") |>
  dplyr::mutate(
    deg_status=factor(deg_status, levels=c("Up", "Down")),
    tissue=ifelse(tissue == "fb_status", "Fat Body", "Whole Body"),
    tissue=factor(tissue, levels=c("Whole Body", "Fat Body"))
  ) |>
  ggplot(aes(x=tissue, y=n, fill=deg_status)) +
    geom_col(width=0.5) +
    theme_bw() +
    theme(
      axis.text.x=element_text(angle=45, hjust=1),
      legend.position="top",
      plot.title.position="plot",
      axis.title.x=element_blank()
    ) +
    labs(
      y="Gene Count", x="Comparison", fill="Differential\nStatus:",
      title="Differentially Expressed Genes\nWith Significant Interaction"
    ) +
    scale_fill_manual(values=c("Down"="salmon", "Up"="darkturquoise"))

print(p)
```

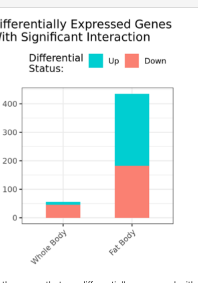


For the genes that are differentially expressed with an interaction term, an illustrative heatmap is generated showing the $log2\,\mathrm{FC}$ for each gene in the whole body and fat body comparisons. The genes are split based on their combined differential status in either of the comparisons, i.e. whether they were up or down-regulated in the whole body and/or fat body.

```r
mat_df <- degs_and_interactions |>
  dplyr::filter(
    (fb_svalue < 0.005 | wb_svalue < 0.005) & interaction_padj < 0.05
  ) |>
  dplyr::mutate(
    group=str_glue(
      "WB ({wb}) & FB ({fb})", wb=wb_status, fb=fb_status
    ),
    group=fct_infreq(group)
  ) |>
  dplyr::select(
    ensembl_gene_id, fb_log2FoldChange, wb_log2FoldChange, group
  ) |>
  dplyr::rename(
    `Fat Body`="fb_log2FoldChange", `Whole Body`="wb_log2FoldChange"
  )

mat <- mat_df |>
  dplyr::select(!group) |>
  column_to_rownames("ensembl_gene_id") |>
  as.matrix()

mat <- mat[, c("Whole Body", "Fat Body")]

p <- Heatmap(
  mat, show_row_names=FALSE, show_column_dend=FALSE, name="log2\nFold Change",
  show_row_dend=TRUE,
  row_split=mat_df[["group"]], row_title_rot=0, row_title_side="left",
  row_title_gp=gpar(fontsize=0), cluster_row_slices=FALSE,
  cluster_columns=FALSE
)

print(p)
```

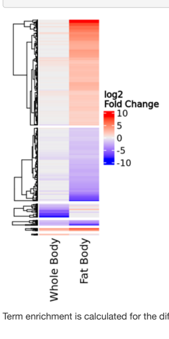


Term enrichment is calculated for the differentially expressed genes with an interaction term.

```r
fb_int <- degs_and_interactions |>
  dplyr::filter(fb_svalue < 0.005 & interaction_padj < 0.05) |>
  dplyr::mutate(group=ifelse(
    fb_log2FoldChange > 0, "Fat Body Up", "Fat Body Down")
  ) |>
  split(~ group) |>
  map(pull, ensembl_gene_id)

wb_int <- degs_and_interactions |>
  dplyr::filter(wb_svalue < 0.005 & interaction_padj < 0.05) |>
  dplyr::mutate(group=ifelse(
    wb_log2FoldChange > 0, "Whole Body Up", "Whole Body Down")
  ) |>
  split(~ group) |>
  map(pull, ensembl_gene_id)

wb_fb_int <- c(wb_int, fb_int)

enrich_results <- list()

enrich_results[["Hallmark Gene Sets"]] <- map(
  wb_fb_int, term_enrichment,
  rownames(dds), category="H", subcategory=""
)

enrich_results[["Reactome Pathways"]] <- map(
  wb_fb_int, term_enrichment,
  rownames(dds), category="C2", subcategory="CP:REACTOME"
)

enrich_results[["Chemical and Genetic Perturbations"]] <- map(
  wb_fb_int, term_enrichment,
  rownames(dds), category="C2", subcategory="CGP"
)

enrich_results[["Gene Ontology Molecular Function"]] <- map(
  wb_fb_int, term_enrichment,
  rownames(dds), category="C5", subcategory="GO:MF"
)

enrich_results[["Gene Ontology Biological Process"]] <- map(
  wb_fb_int, term_enrichment,
  rownames(dds), category="C5", subcategory="GO:BP"
)

enrich_results[["Human Phenotype Ontology"]] <- map(
  wb_fb_int, term_enrichment,
  rownames(dds), category="C5", subcategory="HPO"
)
```

Finally, the pathway enrichment is plotted. The same criteria are used as the first enrichment plot from earlier in the analysis.

```r
p <- imap(enrich_results, plot_enrichment)
p <- wrap_plots(p, ncol=1)

print(p)
```

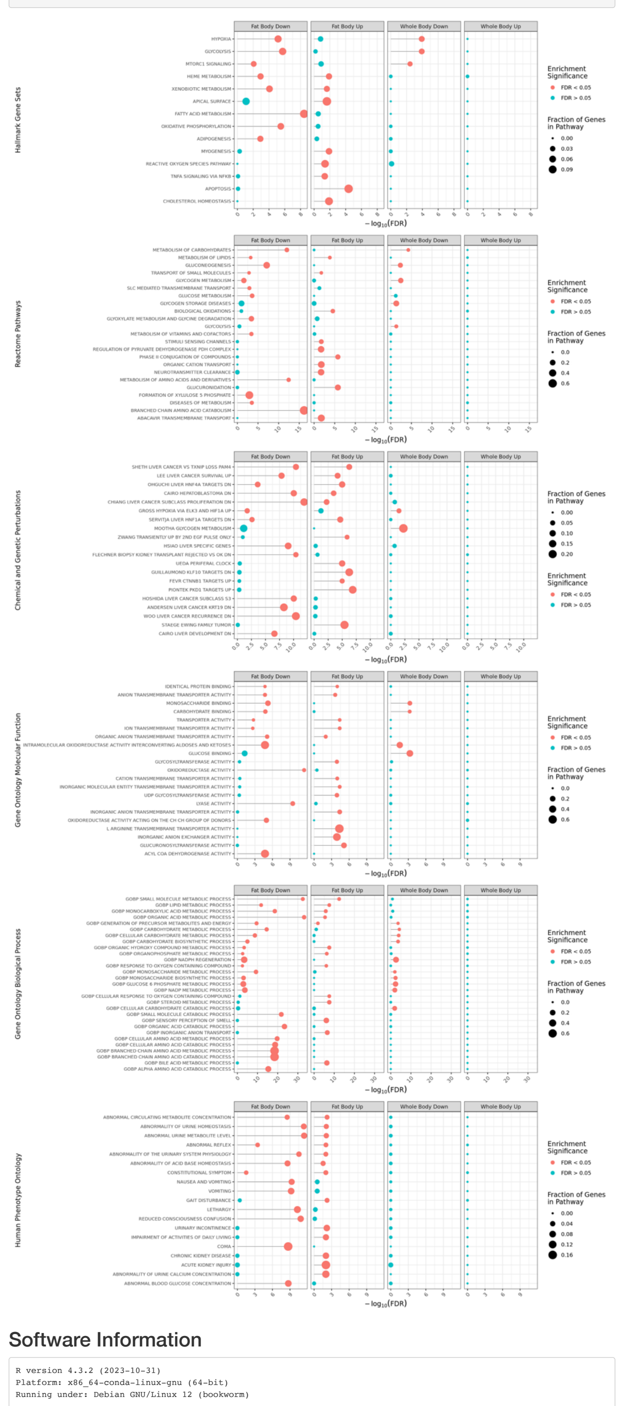

#### Software Information

```
R version 4.3.2 (2023-10-31)
Platform: x86_64-conda-linux-gnu (64-bit)
Running under: Debian GNU/Linux 12 (bookworm)

Matrix products: default
BLAS/LAPACK: /opt/conda/envs/tennessen-rnaseq-err/lib/libopenblasp-r0.3.26.so;  LAPACK version 3.12.0

locale:
 [1] LC_CTYPE=C.UTF-8       LC_NUMERIC=C           LC_TIME=C.UTF-8
 [4] LC_COLLATE=C.UTF-8     LC_MONETARY=C.UTF-8    LC_MESSAGES=C.UTF-8
 [7] LC_PAPER=C.UTF-8       LC_NAME=C              LC_ADDRESS=C
[10] LC_TELEPHONE=C         LC_MEASUREMENT=C.UTF-8 LC_IDENTIFICATION=C

time zone: Etc/UTC
tzcode source: system (glibc)

attached base packages:
[1] grid      stats4    stats     graphics  grDevices utils     datasets
[8] methods   base

other attached packages:
 [1] patchwork_1.2.0             eulerr_7.0.0
 [3] msigdbr_7.5.1               fgsea_1.28.0
 [5] ComplexHeatmap_2.18.0       PCAtools_2.14.0
 [7] ggrepel_0.9.5               DESeq2_1.42.0
 [9] SummarizedExperiment_1.32.0 Biobase_2.62.0
[11] MatrixGenerics_1.14.0       matrixStats_1.2.0
[13] GenomicRanges_1.54.1        GenomeInfoDb_1.38.1
[15] IRanges_2.36.0              S4Vectors_0.40.2
[17] BiocGenerics_0.48.1         biomaRt_2.58.0
[19] tximport_1.30.0             lubridate_1.9.3
[21] forcats_1.0.0               stringr_1.5.1
[23] dplyr_1.1.4                 purrr_1.0.2
[25] readr_2.1.5                 tidyr_1.3.1
[27] tibble_3.2.1                ggplot2_3.4.4
[29] tidyverse_2.0.0

loaded via a namespace (and not attached):
  [1] RColorBrewer_1.1-3         jsonlite_1.8.8
  [3] shape_1.4.6                magrittr_2.0.3
  [5] farver_2.1.1               rmarkdown_2.25
  [7] GlobalOptions_0.1.2        zlibbioc_1.48.0
  [9] vctrs_0.6.5                memoise_2.0.1
 [11] DelayedMatrixStats_1.24.0 RCurl_1.98-1.14
 [13] htmltools_0.5.7            S4Arrays_1.2.0
 [15] progress_1.2.3             curl_5.1.0
 [17] SparseArray_1.2.2          sass_0.4.8
 [19] bslib_0.6.1                plyr_1.8.9
 [21] cachem_1.0.8               lifecycle_1.0.4
 [23] iterators_1.0.14           pkgconfig_2.0.3
 [25] rsvd_1.0.5                 Matrix_1.6-5
 [27] R6_2.5.1                   fastmap_1.1.1
 [29] GenomeInfoDbData_1.2.11    clue_0.3-65
 [31] numDeriv_2016.8-1.1        digest_0.6.34
 [33] colorspace_2.1-0           AnnotationDbi_1.64.1
 [35] dqrng_0.3.2                irlba_2.3.5.1
 [37] RSQLite_2.3.4              beachmat_2.18.0
 [39] labeling_0.4.3             filelock_1.0.3
 [41] fansi_1.0.6                timechange_0.3.0
 [43] polyclip_1.10-6            httr_1.4.7
 [45] abind_1.4-5                compiler_4.3.2
 [47] bit64_4.0.5                withr_3.0.0
 [49] doParallel_1.0.17          BiocParallel_1.36.0
 [51] DBI_1.2.1                  highr_0.10
 [53] MASS_7.3-60                rappdirs_0.3.3
 [55] DelayedArray_0.28.0        rjson_0.2.21
 [57] tools_4.3.2                glue_1.7.0
 [59] polylabelr_0.2.0           cluster_2.1.6
 [61] reshape2_1.4.4             generics_0.1.3
 [63] gtable_0.3.4               tzdb_0.4.0
 [65] data.table_1.14.10         hms_1.1.3
```

```
 [67] BiocSingular_1.18.0       ScaledMatrix_1.10.0
 [69] xml2_1.3.6                utf8_1.2.4
 [71] XVector_0.42.0            foreach_1.5.2
 [73] pillar_1.9.0              emdbook_1.3.13
 [75] babelgene_22.9            vroom_1.6.5
 [77] circlize_0.4.15           BiocFileCache_2.10.1
 [79] lattice_0.22-5            bit_4.0.5
 [81] tidyselect_1.2.0          locfit_1.5-9.8
 [83] Biostrings_2.70.1         knitr_1.45
 [85] xfun_0.41                 stringi_1.8.3
 [87] yaml_2.3.8                evaluate_0.23
 [89] codetools_0.2-19          bbmle_1.0.25.1
 [91] cli_3.6.2                 munsell_0.5.0
 [93] jquerylib_0.1.4           Rcpp_1.0.12
 [95] dbplyr_2.4.0              coda_0.19-4.1
 [97] png_0.1-8                 bdsmatrix_1.3-6
 [99] XML_3.99-0.16.1           parallel_4.3.2
[101] blob_1.2.4                prettyunits_1.2.0
[103] sparseMatrixStats_1.14.0  bitops_1.0-7
[105] mvtnorm_1.2-4             apeglm_1.24.0
[107] scales_1.3.0              crayon_1.5.2
[109] GetoptLong_1.0.5          rlang_1.1.3
[111] cowplot_1.1.3             fastmatch_1.1-4
[113] KEGGREST_1.42.0
```
